# Perirhinal cortex structure and function is dysregulated by corticosteroid treatment

**DOI:** 10.1101/2025.06.06.658287

**Authors:** Matthew D. B. Claydon, Olly Troy, Matthew T. Birnie, Yvonne M. Kershaw, Gareth R. I. Barker, E. Clea Warburton, Zuner. A. Bortolotto, Stafford L. Lightman, Becky L. Conway-Campbell

## Abstract

Glucocorticoids play a crucial role in the stress response and in the regulation of circadian function, as well as cognitive, cardiovascular, metabolic and immunological processes. While synthetic glucocorticoids (sGC) are widely used in treating inflammatory disorders, their impact on cognitive functions, which include memory deficits and dysregulation of mood, remains less understood. Here, we demonstrate that chronic treatment with the sGC methylprednisolone (MPL) dysregulates the synaptic proteome and impairs cognitive function in the perirhinal cortex, a region critical for recognition memory and visual perception. Further, we show that synaptic structure and plasticity are altered by MPL treatment, highlighting the mechanisms through which sGCs disrupt mnemonic processing. These results may have broad implications for understanding the cognitive side effects of widely used sGC treatments.

## Introduction

Glucocorticoids are adrenal steroid hormones that are pivotal in the stress response, but also in the regulation of an array of physiological processes, including metabolism, immune responses, and arousal[1–4]. Synthetic glucocorticoids (sGCs) such as methylprednisolone (MPL) are clinically useful in the treatment of immune disorders and inflammation[5], however, they have significantly different pharmacokinetic and receptor-binding profiles compared to their endogenous counterparts. Treatment with sGCs can result in alterations to endogenous glucocorticoid signaling which can in turn result in serious side effects. Indeed, elevated levels of cortisol (the primary glucocorticoid in humans) or exogenous steroid hormones have been associated with several neuropsychiatric disorders and even psychosis[6–8]. Despite the profundity and prevalence of sGC treatment side-effects, the mechanisms by which these occur in the brain are not fully understood, hampering improvement in the delivery of treatments using these powerful anti-inflammatory drugs.

We have previously reported that disrupting endogenous corticosteroid signaling with MPL treatment results in the dysregulated of hippocampal gene expression, particularly genes associated with synaptic pathways, as well as deficits in synaptic plasticity and cognition[9]. Similarly, the effects of chronic stress and sGC treatment on neuronal excitability and plasticity in the hippocampus and amygdala have been demonstrated[10–14], but how glucocorticoid signaling regulates other medial temporal lobe structures, including the perirhinal cortex (PRh) is significantly less well understood.

The PRh is a region situated in the medial temporal lobe that is best known for its roles in visual perception[15] and the processing of mnemonic information[16]. The pathway from perirhinal to entorhinal cortex (EC) is one of two major entrances for sensory information into the entorhinal-hippocampal network[17] and due to the high level of intrinsic inhibition within the network, it has been posited that the PRh acts to gate information in to the entorhinal-hippocampal network[18, 19]. Interestingly, the PRh possesses neither the columnar organization that admits 6 cell layers of neocortex to process information in a concentrated microcircuit, nor the distributed networks of hippocampus that permit wide ranges of information to be collated in contextual processing[20]. However, PRh neurons are known to receive inputs from many sensory modalities[17] and are able to function as a site of integration, thereby playing an important role in learning and memory[21]. Recognition memory, particularly single item recognition, critically depends on the PRh[22] and neurons in the PRh can be differentially activated by familiar and novel stimuli[23].

Despite its proximity and functional relation to the entorhinal-hippocampal network, which are known to be highly regulated by glucocorticoids, the effects of endogenous and synthetic glucocorticoids on the neurobiology of the perirhinal cortex, an area critical for recognition memory, remain largely unexplored.

## Results

### Methylprednisolone dysregulates the perirhinal proteome

The pulsatile nature of endogenous glucocorticoid release is important in maintaining phasic glucocorticoid receptor (GR) functionality, such as responsiveness to stress-induced glucocorticoid spikes[24], transcriptional regulation in target tissues[25], and even in normal cognitive and emotional responses[26]. Because MPL treatment is known to disrupt the rhythmic expression of RNA in rat hippocampus[9], we first sought to establish the effects of the same course of MPL treatment ((1 mg/mL) for 5-7 days in drinking water (Figure 1A)) on clock gene expression in perirhinal cortex.

**Figure 1.**
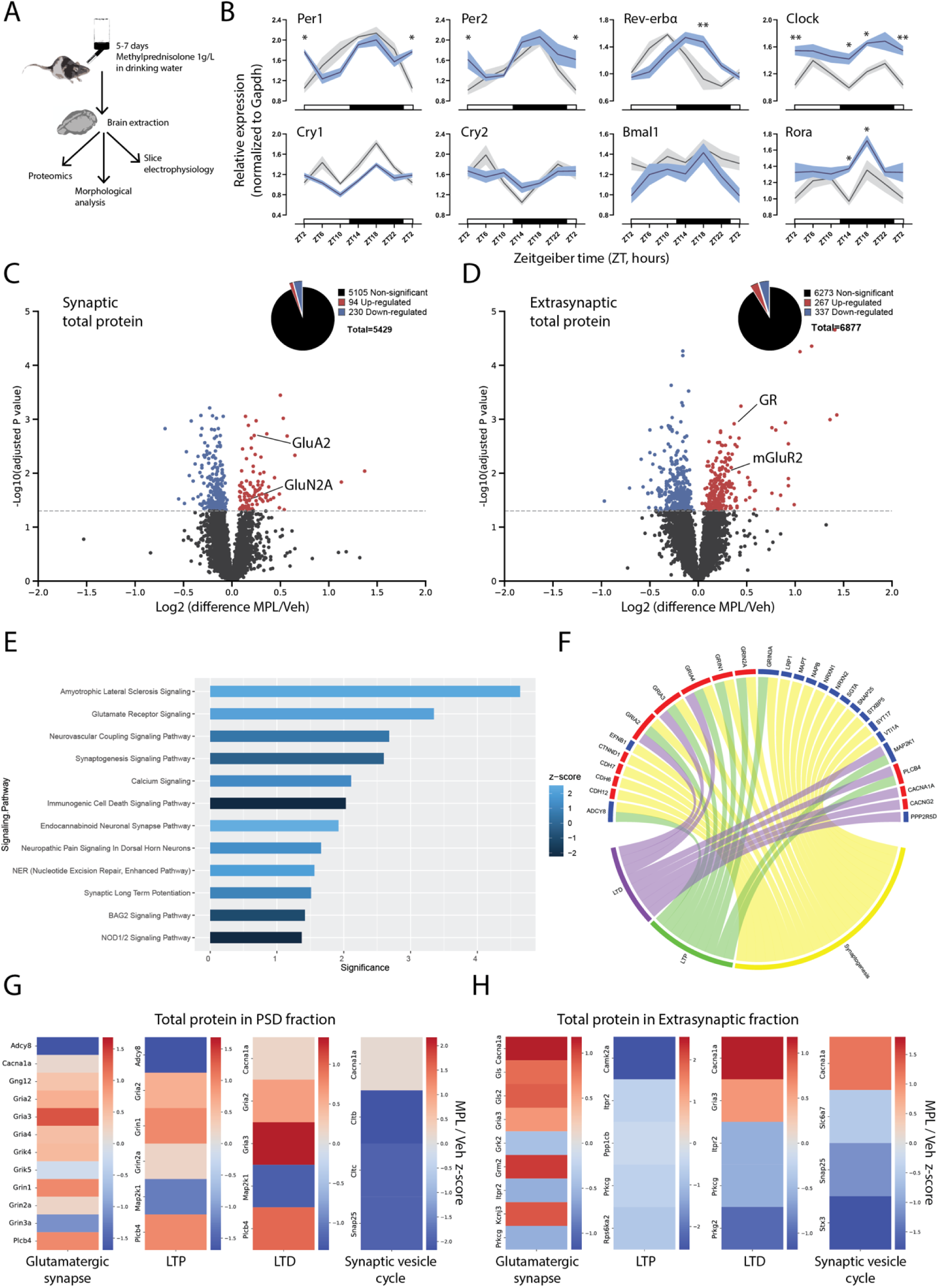
– MPL treatment alters the proteome of synaptic and extrasynaptic fractions from perirhinal cortex. (A) Schematic of experimental timeline. MPL (1 mg/mL) was administered in drinking water for 5-7 days, after which brains were taken or behavioral experiments performed. (B) MPL treatment resulted in the dysregulation of clock gene expression. Two-way ANOVA with Šídák’s multiple comparisons to test for significance at each time point between vehicle (grey) and MPL (blue) treated groups. * = P<0.05, ** = P<0.01. n = 4 rats per treatment group. (C and D) Tandem mass tag (TMT) nanoLC-MS/MS-based proteomic analysis of synaptic/extrasynaptic fractions, respectively. Red represents a significant increase in the MPL group vs. CTL, and blue represents a significant decrease. Significance threshold P<0.05. (E and F) Ingenuity pathway analysis revealed significantly (P < 0.05) regulated biological pathways which were associated with differentially expressed proteins. (G and H) The log fold change of proteins found to be significantly differentially expressed between MPL and CTL groups was quantified by z-score (represented by colour) and grouped by signaling pathway of interest according to the Kyoto Encyclopedia of Genes and Genomes (KEGG).

A significant main effect of time was found for all clock genes bar Cry2, indicating that like the hippocampus, the expression of molecular clock machinery in PRh is temporally regulated. Crucially, the oscillations in clock gene expression across the photoperiod were found to be sensitive to treatment with the sGC MPL (Figure1B). Given the established role of circadian rhythms in regulating various aspects of neuronal function, including synaptic plasticity[9], this suggests that glucocorticoid treatment could disrupt the timing and expression of these processes, which are crucial for proper cognitive function. We hypothesized, therefore, that these changes to GR signaling and clock gene expression may therefore result in aberrant protein expression crucial for cognitive function during the active phase, a time critical for the expression of plasticity associated genes[9] and learning and memory processes.

To address this, we utilized tandem mass tag (TMT) nanoLC-MS/MS-based proteomic analysis of perirhinal tissue isolated at the onset of the active phase from rats treated with either MPL or vehicle. Proteomic analysis of synaptic fractions revealed that out of 5,429 proteins analyzed, 324 (5.97%) were differentially expressed (Figure 1C). In the extra-synaptic fraction, 604 out of 6,867 proteins (8.65%) were differentially expressed in the perirhinal cortex of MPL-treated rats (Figure 1D). The extensive impact of MPL treatment on synaptic and extra-synaptic proteomes suggest that there may be systemic alterations to specific signaling pathways that contribute to sGC induced cognitive deficits.

Ingenuity Pathway Analysis (IPA) was employed to identify neurobiological processes that were associated with differentially expressed proteins (DEPs) (Figs 1E & F). Notably, alterations in synaptic pathways, which are well-documented in the context of corticosteroid-induced cognitive deficits, became a central focus of the analysis, and protein changes between conditions were quantified according to signaling pathway.

Several key proteins associated with synaptic plasticity, including the AMPA and NMDA receptor subunits *Gria2*, *Gria3*, *Grin2a*, and *Grin3a* were found to be differentially expressed in the synaptic fraction. Proteins crucial in the exocytosis of synaptic vesicles such as *Snap25* were also found to be significantly downregulated following MPL treatment (Figs 1G & H).

In the extrasynaptic fraction, synaptic signaling proteins were also found to be differentially expressed, including *Grm2* and *Gria3.* Interestingly, *Camk2a,* a signaling protein essential in the formation of LTP, was found to be highly downregulated by MPL treatment.

The positions and abundances of phosphorylation residues were also analyzed alongside the total protein data in each sample, with 8423 phosphopeptide sites identified in the PSD fraction and alongside 8792 in the extrasynaptic fraction. In the PSD fraction, significant alterations to the phosphorylation state of 245 proteins were identified (Figure 2A), including critical proteins such as *Camk2a* and *Grin2b* (Figure 2B) which have both have been shown to play central roles in the stress-mediated changes to hippocampal function[27, 28]. Indeed, the increased phosphorylation at S331 observed here has previously been shown to be associated with inhibition of CaMKIIa activity[29]. It has been hypothesized that MPL induced synaptic deficits in hippocampus are mediated by the disruption of CaMKIIa and GluN2B-NMDAR interactions[9], and strikingly in the perirhinal cortex we found an increase in the phosphorylation of *Camk2a* T337, which has been shown to occlude the interaction of CaMKII with GluN2B, a process which is pivotal in the expression of synaptic plasticity[30].

**Figure 2.**
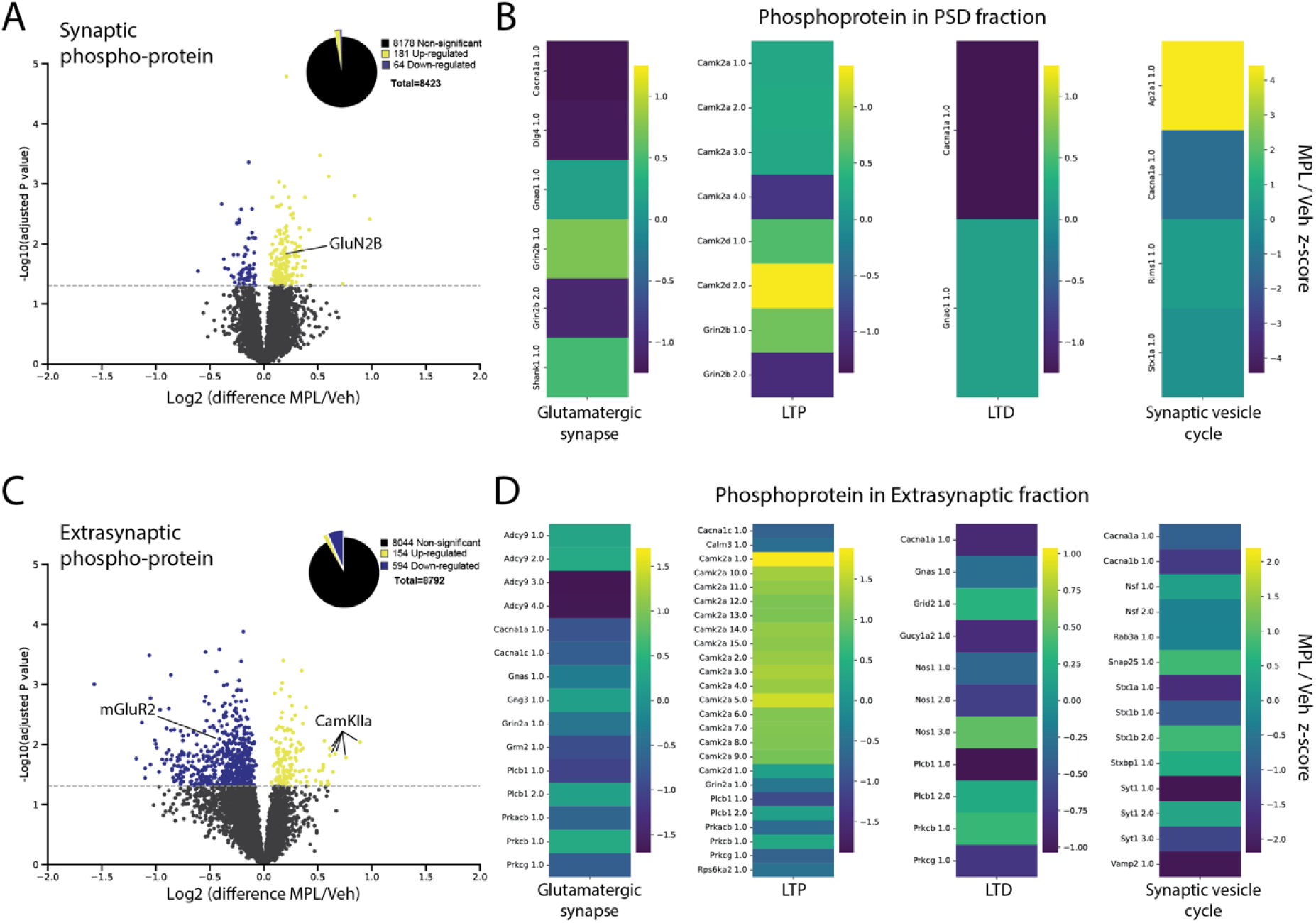
– Methylprednisolone induced changes to perirhinal phosphoprotein. (A and C) Tandem mass tag (TMT) nanoLC-MS/MS-based proteomics revealed changes to the phosphorylation state of many proteins in the post-synaptic density (B) and extra-synaptic fraction (D). Significantly regulated proteins (–log10 (0.05)) are highlighted in colour. Downregulation and upregulation by MPL of phosphorylation are indicated by negative and positive values on the x axis, respectively. (B and D) Proteins with altered phospho-sites were sorted by KEGG pathway analysis, and significantly regulated phospho-sites quantified by Z scoring log fold change.

In the extrasynaptic fraction 748 phosphopeptide sites exhibited significant changes (Figure 2C) including *Grin2a*, *CamKIIa*, and *Mapt*, further implicating NMDA receptor and CaMKII dysregulation in the deleterious effects of MPL treatment. These data suggest that sGCs may induce cognitive deficits not only by influencing protein expression, but also by modulating the functional state of these proteins through phosphorylation changes, a process known to play a critical role in regulating synaptic strength and plasticity[31].

### MPL treatment decreases spontaneous excitatory neurotransmission onto perirhinal neurons

As expression of proteins associated with the synaptic vesicle cycle and their phospho-states were altered by MPL treatment, we sought to investigate whether this was reflected in functional deficits to synaptic transmission. To do so, we recorded miniature excitatory and inhibitory post synaptic currents (mEPSCs and mIPSCs) from perirhinal cortex pyramidal cells. Strikingly, MPL treatment resulted in a decrease in the frequency, but not amplitude, of mEPSCs (2.01 ± 0.22 vs 1.10 ± 0.15 Hz, Figure3B and C). This decrease in frequency could reflect a reduction in synaptic release probability, or a reduction in the number of functional excitatory synapses.

Interestingly, MPL treatment did not result in any changes to mIPSC properties (Figure3E and F), suggesting that chronic steroid treatment may preferentially dysregulate excitatory neurotransmission in the perirhinal cortex. Indeed, MPL induced changes in the expression of a greater number of proteins associated glutamatergic synapses than GABAergic (Figure1K). These data suggest that MPL treatment disrupts the balance between excitatory and inhibitory (E-I) input to perirhinal neurons, which is thought to be crucial in the tuning of circuits to respond appropriately to salient stimuli and the efficient coding of information[32, 33]. Importantly, cortical E-I balance has also been shown to be dysregulated in autism spectrum disorder and is frequently altered in various neuropsychiatric disorders[34].

Whole-cell patch clamp experiments (Figure 3G) confirmed that MPL induced alterations to the expression of AMPA and NMDA receptor subunits observed in the proteomics data were reflected in changes to the functional activity of AMPA and NMDA receptors in perirhinal cortex neurons, as measured by AMPA:NMDA ratios (Figure 3H). However, no significant effect on the GluN2A/2B subunit composition of NMDA receptors was observed (Figure 3I).

**Figure 3.**
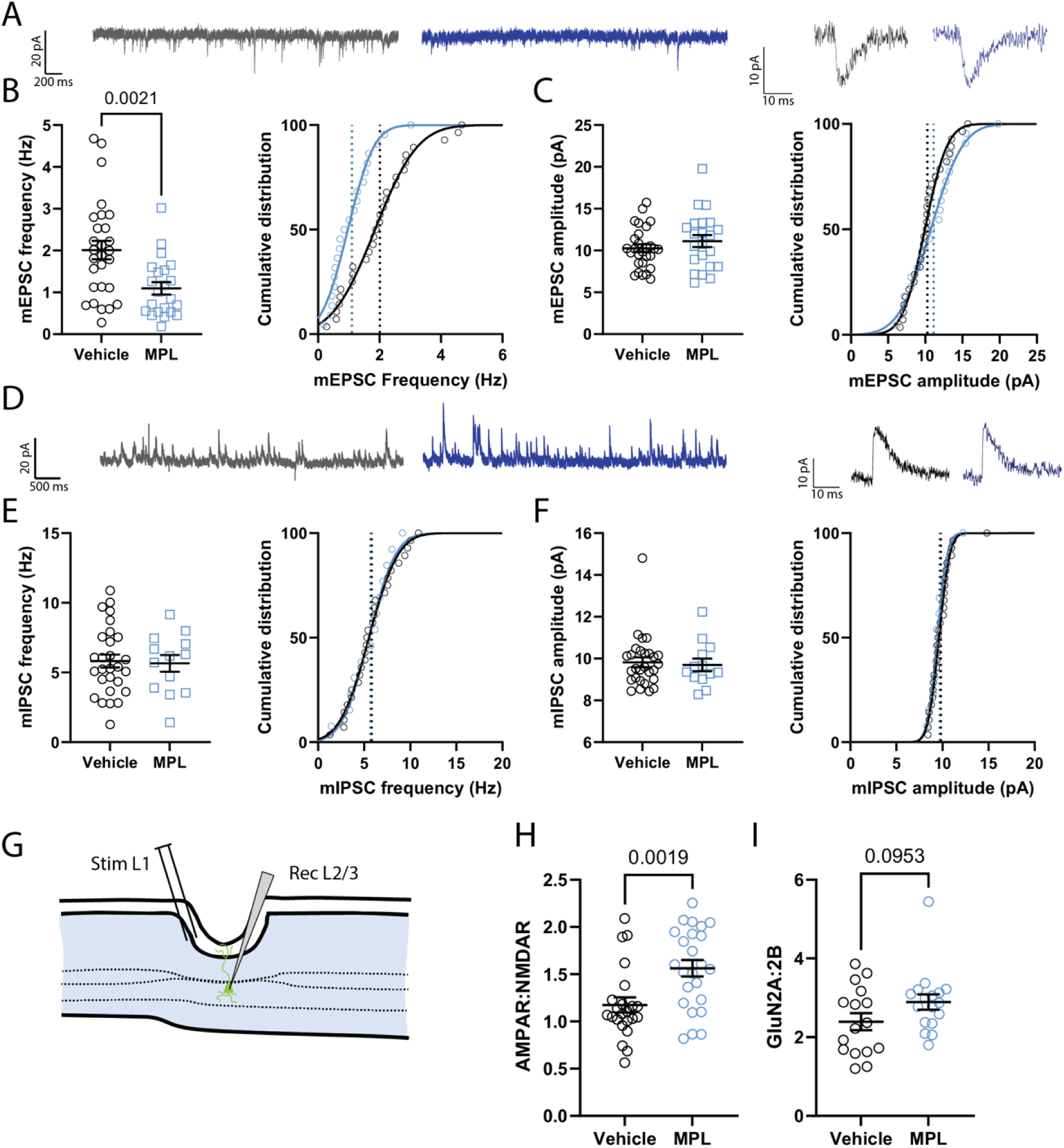
– The frequency of spontaneous excitatory neurotransmission is reduced by MPL treatment. (A) Representative traces of mEPSC recordings (left) and individual events (right) from vehicle (grey) and MPL (blue) treated rats. (B) MPL treated induced a significant reduction in the frequency but not amplitude (C), of mEPSC events in perirhinal cortex neurons (n = 28, 8 [Veh], and 22, 8 [MPL]) (n = cells, animals) (P < 0.01). (D) Representative traces of mIPSC recordings from vehicle and MPL treated rats. (E and F) MPL treatment did not alter mIPSC frequency or amplitude in perirhinal cortex (n = 28, 8 [Veh] and 13, 4 [MPL]) (n = cells, animals). (G) Neurons in layer II/III of perirhinal cortex were accessed in whole-cell patch clamp and excitatory post-synaptic potentials (EPSCs) evoked by stimulating in layer I. (H) The ratio of AMPA:NMDA mediated currents in layer II/III PRh neurons was significantly increased in MPL treated rats compared to vehicle treated controls (n = 23, 12 [Veh] and 24, 12 [MPL]) (n = cells, animals) (P < 0.01). (I) GluN2A:GluN2B ratio measured by peak current at baseline vs. following 20 minutes application of 5 μM Ro-25-6981 was not significantly altered. Unpaired t-tests were used to test for significance and cells treated as experimental units following testing with a linear mixed model to assess sources of variance.

### Morphological changes to dendrites and spines

The alterations to the synaptogenesis associated proteome and reductions in mEPSC frequency observed suggest that MPL treatment may alter neuronal morphology to decrease the number and functionality of synapses in perirhinal cortex. Indeed, glucocorticoid hormones are known regulators of neuronal morphology in the hippocampus and chronic stress has been shown to alter the morphology of perirhinal dendritic spines in a CRH dependent manner.

Therefore, we utilized the Golgi-cox staining method to investigate how MPL treatment may affect the structural morphology of PRh pyramidal cells. Surprisingly, Sholl analysis revealed an MPL-induced increase in the complexity of the proximal apical dendrites (<150 µm from soma), specifically at 40-60 µm radial distance from the soma, while distal apical and basal dendrites were unaffected.

Moreover, no changes to spine density in L5 PRh pyramidal cells could be observed following MPL treatment in any dendritic region (Figure 4C).

**Figure 4.**
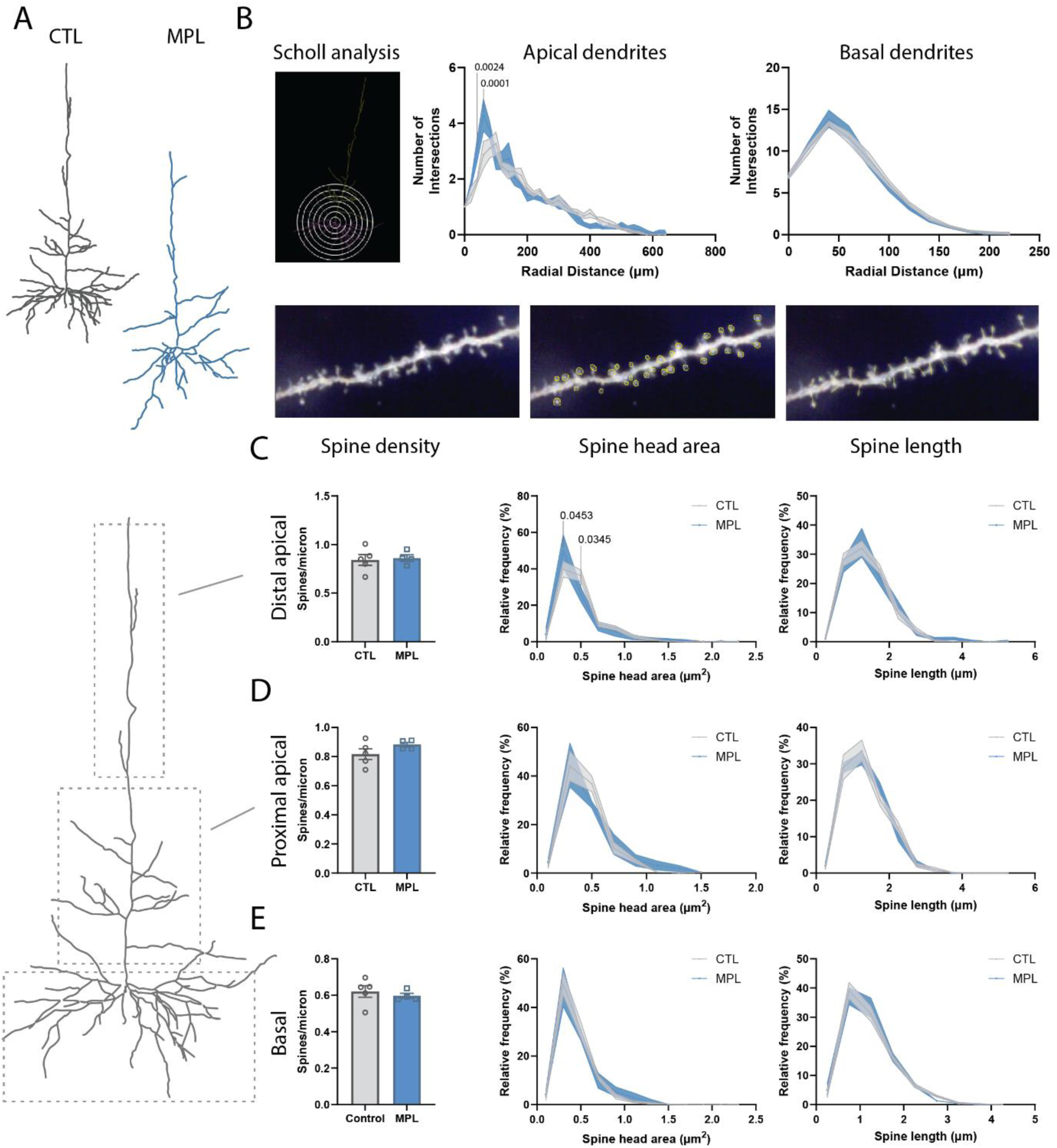
– Dendritic and spine morphology of perirhinal cortex principal cells is altered by MPL treatment. (A) Representative traced neurons from Golgi-cox stained slices from vehicle treated (grey, n = 5) and MPL treated (blue, n = 4) rats. (B) Sholl analysis revealed a significant effect of the interaction between drug treatment and radial distance on the complexity of apical dendrites of perirhinal neurons in rats treated with MPL compared to vehicle (Two-way ANOVA, F_32, 231_ = 2.215, P=0.0004, Šídák’s multiple comparisons test). (C, D, and E) Spine density, spine head area, and spine length were calculated for the distal apical (> 150 µm from soma), proximal apical (< 150 µm from soma), and basal dendritic compartments. n = 4 (veh) and 5 (MPL). Two-way ANOVA with Šídák’s multiple comparisons test were used to test for statistical significance and animals treated as experimental units.

However, MPL treatment may have induced subtle changes to the relative frequency of smaller and larger dendritic spines. Despite no significant main effect of treatment or interaction between treatment and spine head area (interaction P=0.0796), a significant increase in the relative frequency of spines with a spine head area of 0.3 µm following MPL treatment (49.87 ± 8.96 vs. 39.77 ± 4.35 %, P = 0.0453), and a decrease in the frequency of spines with a larger spine head area of 0.5 µm (25.96 ± 3.95 vs. 36.38 ± 3.13 %, P = 0.0345) (Figure 4C). No significant effects on spine head area or length were seen in any other dendritic region (Figure 4D and E). This is in corroboration with previous findings that dendritic spines have been shown to respond differently to stress depending on distance from soma and location in either basal or apical sites. Indeed, in rat cortex, distal spines were most susceptible to decreases in spine volume and surface area following repeated stress [35] while spine density was only affected in apical and not basal compartments [36]. These data indicate that alterations to mEPSC frequency may reflect a reduction in presynaptic release, which is corroborated by the alterations to expression of proteins associated with the synaptic vesicle cycle (i.e. downregulation of SNAP25, and the clathrin light and heavy chain encoding proteins Cltb and Cltc, respectively). The shift in spine head area to an increased frequency of spines with reduced spine head area may also reflect the early stages of structural changes, leading to fewer functional synapses in time.

### Methylprednisolone treatment induces deficits in perirhinal synaptic plasticity

In the post-synaptic density (PSD) fraction, DEPs included the GluA2 subunit of the AMPA receptor and the GluN1 and GluN2A subunits of the NMDA receptor (Figure 1G), which are crucial in glutamatergic transmission and the expression of synaptic plasticity [37, 38]. Their roles are especially prominent in long-term depression, the primary plasticity form in the PRh; GluA2 internalization is a key mechanism in both NMDA and mGluR-dependent LTD[39–41], whilst overexpression of GluN2A has been demonstrated to impair LTD in both the lateral amygdala and the hippocampus[42]. We therefore hypothesized that MPL-induced alterations to glutamatergic receptor subunit expression in the PSD of perirhinal neurons, and changes to the phospho-states of key plasticity proteins, would culminate in deficits to synaptic plasticity. NMDA receptor dependent LTD in the perirhinal cortex is a physiologically relevant paradigm crucial for the expression of recognition memory in rodents[43], and so we sought to test whether LTD induced by 10 minutes of 5 Hz stimulation was disrupted by MPL treatment.

Ex-vivo brain slices containing perirhinal cortex were prepared from rats treated with MPL or vehicle, and electrical stimulation was used to elicit field excitatory post-synaptic potentials (fEPSPs) (Figure 5A). LTD of fEPSPs was reliably induced in the test pathway of slices from vehicle (85.79 ± 1.398 % of baseline, Figure 5B), but not MPL (100.9 ± 2.087 % of baseline, Figure 5C), treated rats.

It has been suggested that GluN1/GluN2A and GluN1/GluN2B NMDA receptors may be preferentially involved in the induction of LTP and LTD, respectively, due to their differing depolarization kinetics[44]. Administration of the selective GluN2B antagonist Ro-25-6981 to the slice prior to LTD induction revealed that the LTD protocol sensitive to MPL treatment was dependent on GluN2B-containing NMDARs (Figure 5E). These data indicate that the MPL-induced dysregulation of the PRh proteome (Figure 1) results in functional deficits to neuronal plasticity in the perirhinal cortex.

**Figure 5.**
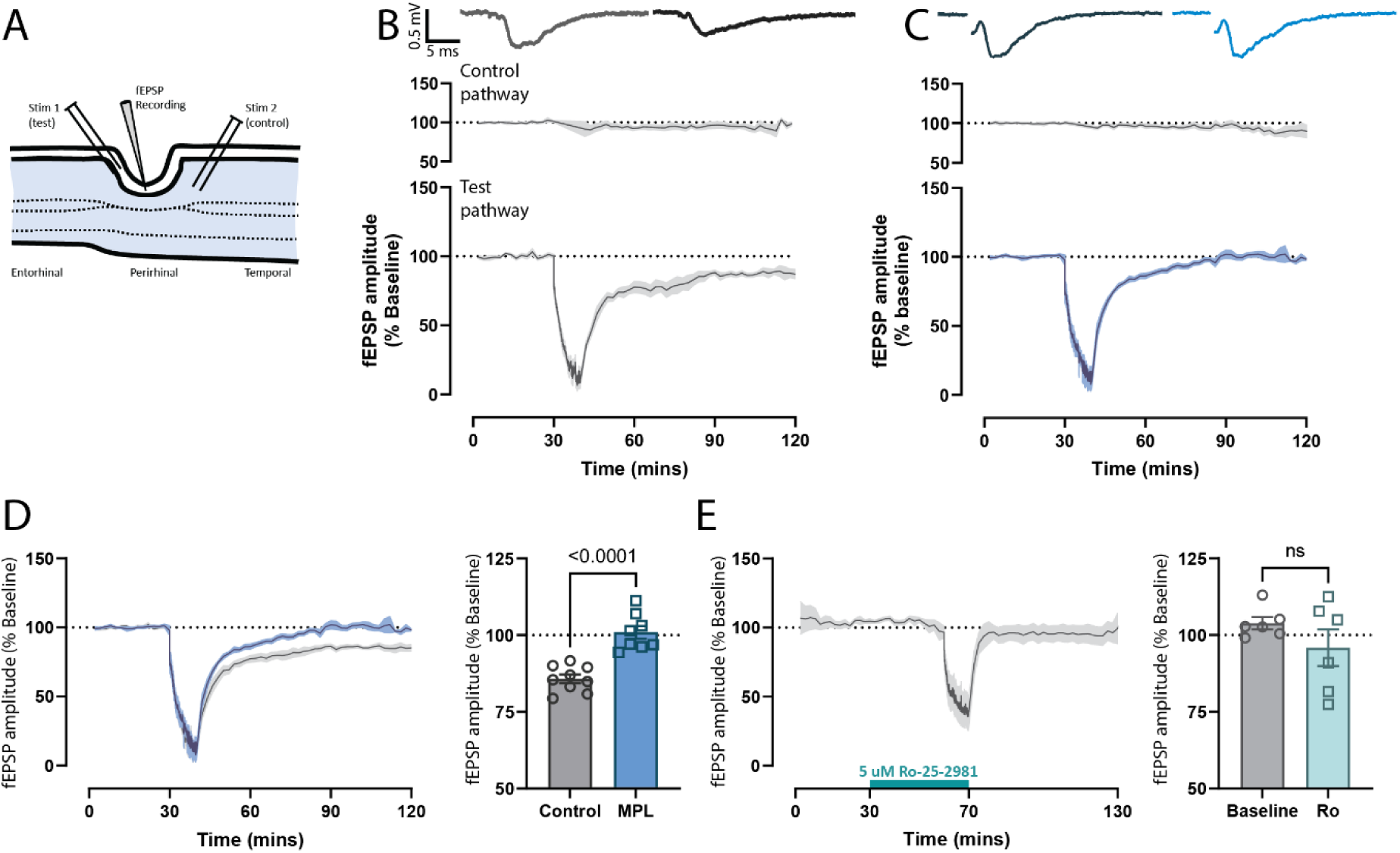
– Long-term depression in perirhinal cortex is disrupted by MPL treatment. (A) Schematic of a perirhinal field recording set up with a control pathway and test pathway stimulating layer 1. The recording electrode was placed at the boundary of layers I/II to record dendritic depolarization of deeper layer neurons. (B) Long-term depression (LTD) of inputs to perirhinal neurons could reliably be induced by 10 minutes of 5 Hz stimulation (n = 9 rats) (C) LTD was not induced by 5 Hz 10 min stimulation in perirhinal cortex slices of rats treated with MPL (n = 8 rats) (D) The degree of LTD induced in perirhinal cortex was significantly reduced in MPL treated animals vs. vehicle treated (unpaired t-test, P < 0.0001) (E) Blockade of GluN2B-containing NMDA receptors with 5 µM Ro-25-6981 for 30 minutes prior to and during the low frequency stimulation (LFS) prevented the induction of LTD

### Neurobiological changes induced by MPL treatment result in deficits in a perirhinal-dependent memory task

Given the MPL induced dysregulation of synaptic protein expression & phosphorylation, synaptic structure, and synaptic plasticity in the perirhinal cortex, we next investigated whether perirhinal cortex dependent behavior was similarly disrupted by MPL treatment.

The NMDA receptor dependent plasticity of PRh synapses that was ablated by MPL treatment is known to be crucial in the expression of single-item recognition[43], and so we hypothesized that this form of memory would be disrupted by MPL treatment. Indeed, in a novel object recognition (NOR) task, retrieval of memory at intermediate (6 hour, 0.218 ± 0.039 vs. –0.070 ± 0.077) and long-term (24 hour, 0.272 ± 0.082 vs. –0.171 ± 0.070) test phases was found to be significantly impaired (Figure 6C and D). Interestingly, NOR memory at a 1-hour interval between sample and test trials was unaffected, indicating that the dysregulation of perirhinal synaptic protein expression and synaptic physiology induced by MPL treatment culminates in ablation of perirhinal dependent long-term, but not short-term, memory.

**Figure 6.**
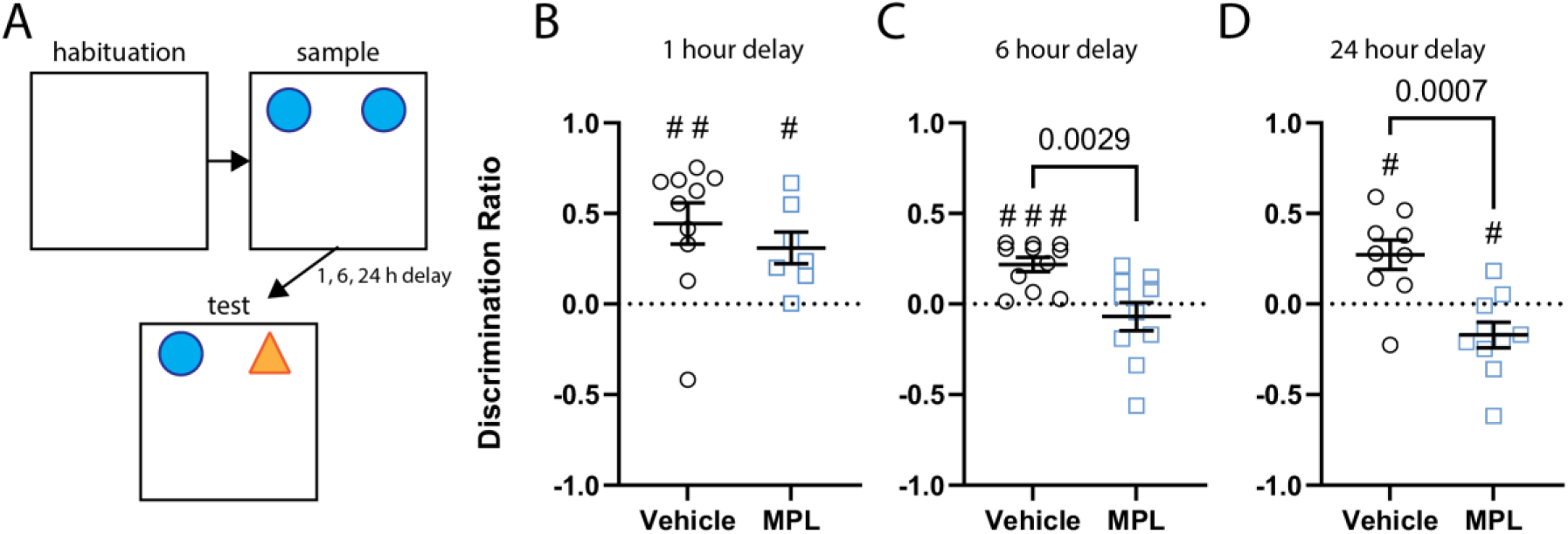
– MPL treatment disrupts long-term novel object recognition memory. (A) Schematic of the novel object recognition (NOR) task. Rats were subject to a 1, 6, or 24 hour delay between the sample and test phase. (B) MPL treatment induced a deficit in the NOR memory task at 6 (n=11, 10) and 24 (n=9, 10), but not 1 (n=10, 7), hours sample/test intervals (# = P < 0.05, ### = P < 0.001, paired t-test vs baseline, unpaired t-test between treatment groups).

## Discussion

Corticosteroids are a major treatment option for a variety of inflammatory and autoimmune diseases, despite a broad side-effect profile including both metabolic and psychiatric disturbances. Psychiatric adverse effects include corticosteroid-induced mania, suicidal thoughts, and cognitive impairment and have been well characterized; despite this, the precise mechanisms mediating them are still poorly understood. It is now apparent that chronic sGC treatment risks can manifest differently to those of chronic stress or endogenous glucocorticoid treatment and a separate body of research is necessary [45].

In the present study, we demonstrate that MPL treatment, designed to mimic a clinical sGC treatment course, leads to myriad dysregulation of perirhinal cortex function: MPL treatment significantly alters the perirhinal proteome, synaptic plasticity and function, neuronal morphology, and ultimately results in diminished long-term memory.

MPL treatment induced potent and broad changes to the perirhinal proteome, which was expected given the extensive genetic regulation mediated by the glucocorticoid and mineralocorticoid receptors[46]. Pathway analysis of DEPs highlighted glutamatergic synapse function, calcium signaling, and synaptic plasticity as key areas warranting further exploration.

Indeed, DEPs in the synaptic fraction included AMPA and NMDA receptor subunits, and the phosphorylation states of GluN2B and CaMKII were found to be altered by MPL in a manner predicted to disrupt their interaction. Given the importance of NMDA receptor activation in perirhinal synaptic plasticity[47, 48] and learning and memory[49, 50], the disruption to PRh LTD observed thereafter was not surprising.

We hypothesize that the differential expression of NMDA receptor subunits and the altered phosphorylation of key residues induced by MPL prevents the association of GluN2B with CaMKII (a protein also regulated by MPL). This protein-protein association is pivotal in the expression of synaptic plasticity[30], and so provides a putative mechanism by which MPL treatment ablates perirhinal LTD. The disruption of this association could result in a reduction in the number of functional NMDA receptors anchored to the synapse, which could account for the increased AMPA:NMDA ratio observed in perirhinal neurons. This notion is supported by the fact that mEPSC amplitude was unchanged, suggesting that AMPA:NMDA increases would be due to a decreased NMDA receptor mediated current. However, the AMPA receptor subunit GluA2 was also found to be significantly upregulated by MPL treatment, and this could also contribute to an increase in AMPA:NMDA. Analysis of the synaptic fraction also revealed a significant reduction in the expression of Adenylyl cyclase 8, which is critical in the regulation of cyclic AMP (cAMP) signalling. Adcy8 knockout studies have demonstrated a key role for this membrane protein in synaptic plasticity in the hippocampus[51, 52], although it’s role in LTD specifically is contentious[53, 54], and may be protocol and region specific. Thus, the high degree of Adcy8 downregulation at the synapse observed here is likely to contribute to deficits in plasticity. Indeed, it seems likely that it is a combinatorial effect of the broad and significant alterations to the synapse associated perirhinal proteome that contributes to a reduction in the expression of physiological synaptic plasticity following treatment with MPL.

The dendritic morphology of both cortical and hippocampal neurons is known to be susceptible to dysregulated HPA axis input. Indeed, chronic stress induces dendritic atrophy in brain regions closely related to the perirhinal cortex[36, 55]. Interestingly, we find here that MPL treatment alters the structure of the dendritic tree differentially, despite producing cognitive impairments in a perirhinal-dependent task. Surprisingly, and in contrast to the hippocampus, MPL-treatment increased the branching proximal to the soma. The reasons for this distinction are unclear, but the biphasic and region-specific nature of many glucocorticoid actions may provide some explanation[56].

Differences in BDNF expression, a key factor in dendritic remodeling in stressful conditions[36, 55], in different brain regions following periods of heightened glucocorticoid levels[57], could also underlie these region-specific effects.

Dendritic spines have a dynamic structure, facilitating the plasticity necessary for complex brain function. However, changes in spine number and structure can be pathological, and a decrease in spine density is a hallmark of neurodegenerative diseases[58]. In the present study, no changes to spine density could be observed in any dendritic region, which was surprising given the reductions in density that have been reported in functionally linked brain regions following chronic stress or glucocorticoid treatment. However, the treatment conditions across these studies vary greatly, and the findings that sGC treatment alone can illicit different effects on spine morphology depending on brain region and treatment course [59, 60] suggest that specific treatment conditions must be carefully considered.

The reduction in spine head area in the distal apical region supports results seen in previous chronic stress studies[42, 61] and suggests that the spine population may be transitioning to a less stable state, predisposing a loss in spine number[59, 60, 62].

Additionally, the dendritic response to stress has been demonstrated to follow distinct temporal dynamics in different brain regions[57], further highlighting the possibility of a gradually changing morphological profile.

We also observed reductions in mEPSC frequency that could reflect a decrease in pre-synaptic release probabilities but also a decrease in the number of active synapses.

Considering this, the reduction in mEPSC frequency may be reflective of the reduction in spine head area observed in the apical dendrites of these cells. This does not preclude that there may also be changes in presynaptic release probability driving a reduction in network activity and resulting in altered spine structure and function. Indeed, proteins involved in the synaptic vesicle cycle and release (SNAP-25, Cltb, and Cltc) were significantly down-regulated by MPL treatment in the synaptic density. SNAP-25 deletion has been shown to reduce the frequency of spontaneous excitatory neurotransmission[63, 64] and spontaneous inhibitory neurotransmission[65], indicating that if the reduction in synaptic SNAP-25 expression is driving the observed decrease in mEPSC frequency, it is only in excitatory terminals. The degree to which the structural changes to spine head width and synaptic vesicle cycle protein expression changes are contributing to the selective reduction in mEPSC frequency, therefore, remains unclear.

In the present study, we observed that deficits in synaptic physiology and NMDA receptor LTD in PRh culminated in the ablation of long-term, but not short-term recognition memory. It has been shown previously that NMDA receptor dependent LTD of PRh synapses is essential for the functional expression of visual recognition memory in rodents[43] (although other receptors are also known to play crucial roles in recognition memory[66, 67]) and although this phenomenon can be observed in many cortical areas along the ventral visual pathway, the endurance of the memory is considerably longer in the PRh than elsewhere[68].Moreover, blockade of perirhinal NMDA receptors was shown to impair the consolidation of long-term novel object recognition memory, while short-term (20 minute delay) memory was unaffected[69, 70], supporting the hypothesis that MPL induced disruption to NMDA receptor dependent LTD underlies the cognitive deficits in NOR memory.

This study used only male rats and thus does not consider the effects of sGC treatment on perirhinal function in females. Interestingly, recent studies have demonstrated that male rats are more susceptible to recognition memory impairment following both acute and chronic stress [71, 72], highlighting the importance of this distinction. Moreover, hippocampal plasticity has been shown to depend on locally synthesized estrogen[73] and blocking estradiol synthesis ablated plasticity in only female animals[73, 74]. While interpretation of the interaction of endogenous and synthetic glucocorticoid hormones with these processes remains difficult, it nonetheless warrants further investigation.

In summary, we have observed a deficit in rat long-term recognition memory following treatment with MPL, a synthetic glucocorticoid commonly prescribed in clinic. This is likely mediated in part by the described deficits in synaptic plasticity in the perirhinal cortex, a brain region that plays a crucial role in recognition memory. These data highlight potential targets for therapeutic intervention, whereby restoring normal signaling could mitigate the cognitive deficits commonly associated with corticosteroid treatment[75]. Deficits in synaptic plasticity processes are linked with the memory impairments observed neurodegenerative disease[76] and as such, impairment of synaptic plasticity in PRh represents a significant risk in the clinical use of MPL. However, advancing our understanding of brain-region and corticosteroid specific effects brings us closer to the prevention of serious neuropsychiatric side effects and the tailoring of treatment regimens for susceptible individuals.

## Methods

All procedures described here were conducted in accordance with the UK Animals (Scientific Procedures) Act, 1986.

## MPL treatment

Methylprednisolone (MPL) as the sodium succinate (Solu-Medrone, Pfizer) was obtained from University Hospitals Bristol pharmacy stores. 1 g of sterile MPL as the sodium succinate was dissolved in 1 L tap water and kept in opaque drinking bottles due to light sensitivity. Normal drinking water was replaced with 1 g/L MPL solution, to which animals had ad-libitum access. Drinking water was then monitored to check consumption rates and weight loss in treated animals was also recorded.

## Trunk blood collection and plasma isolation

During brain slice preparation, immediately following decapitation, the trunk blood was collected from the body of the rat and mixed with 100 μL EDTA on ice to prevent coagulation. Blood samples were then centrifuged for 10 mins at 2,000 x g at 4 °C. Plasma supernatant was then collected in 500 μL aliquots and stored at –20 °C for later analysis.

## Radioimmunoassay for Corticosterone

Using an automated gamma counter (PerkinElmer, US), a CORT radioimmunoassay was carried out to assess CORT levels in blood samples collected from trunk blood. An 11-point standard curve of known CORT concentrations was prepared in B-buffer (25 mM tri-sodium citrate, 50 mM sodium dihydrogen orthophosphate, 1 mg/mL bovine serum albumin: pH3). Plasma obtained either from the automated blood sampler or trunk blood was diluted in triplicate at a ratio of either 1:10 or 1:50 in B-buffer, respectively. A specific CORT antibody (kindly provided by G. Makara, Institute of Experimental Medicine, Budapest, Hungary) was diluted at a ratio of 1:50 in B-buffer and 50 μl added to 100 uL standards, unknown samples, and Quality Control (QC20 and QC100) tubes.

Tracer (Izotop, Institute of Isotopes, Hungary) was diluted in B-buffer to give total counts of 3750cpm in 50μl and added to all tubes (50 μl/tube). Tubes were incubated overnight at 4 °C. Charcoal suspension (5 g charcoal added to 0.5 g Dextran T70 dissolved in 1 L B-buffer) was prepared and 500 μl added to all tubes and briefly vortexed. Blocks were centrifuged at 4000 rpm at 4°C and the resulting supernatant aspirated off. Unknown samples were determined from interpolation of the standard curve.

## Quantitative polymerase chain reaction

Quantitative Polymerase Chain Reaction (qPCR) was used to assess mRNA transcript levels across time points and treatments using ThermoFisher Scientific reagents.

## Sample preparation and primer design

Primers for RNA analysis were designed NCBI BLAST Primer design tool (https://blast.ncbi.nlm.nih.gov/Blast.cgi) or taken from the literature for all Sybr Green assays. Primer length was designed to be between 60-150 bp in length. Nuclease free water was added to each lyophilised primer to give a stock concentration of 100 μM. 20 μL of stock primer was added to 980 μL of nuclease free water, creating a working primer concentration of 2 μM. A standard curve for each primer was produced to assess primer efficiency and to validate the absence of primer dimers. Serial dilutions of cDNA for RNA analysis were prepared to provide a range of cycle threshold (Ct) values that were anticipated would cover the range of values expected to be produced by samples.

**Table 1.1.**
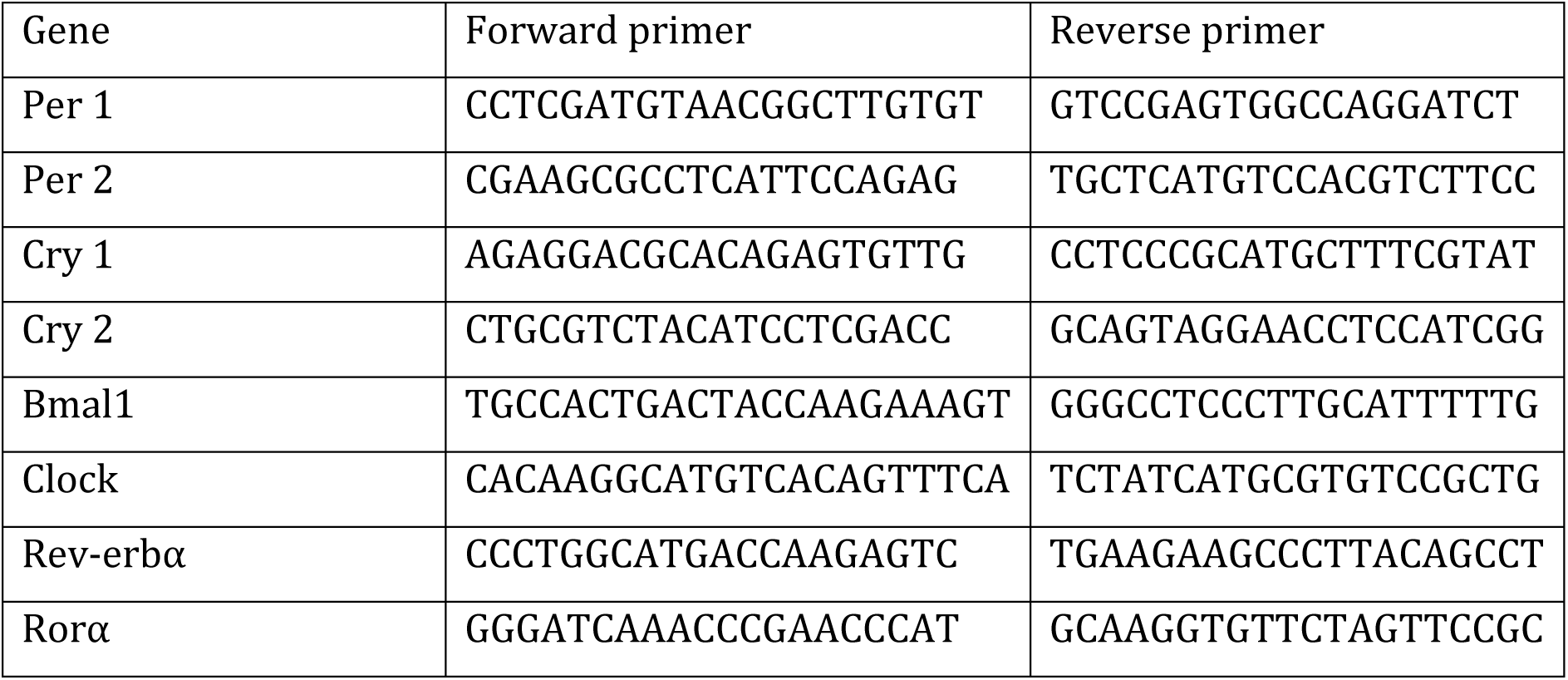
mRNA primer sequences used for Sybr Green qPCR.

**Table 1.2.** Reaction quantities for qPCR.

| Reaction component | Volume per well (μL) |
| --- | --- |
| Fast Sybr®Green Master mix | 5 |
| Forward Primer | 1.5 |
| Reverse Primer | 1.5 |
| Nuclease-free water | 1 |
| cDNA sample | 1 |
| Total Reaction volume | 10 |

## qPCR reaction

qPCR determined the approximate levels of mRNA transcripts in RNA samples. Experiments were performed using 96-well plates (Applied Biosystems, Life Technologies, UK) with a 10 μL final reaction per well (Table 2.2). Each PCR reaction contained 1μL of cDNA. Samples were pipetted in duplicate and a no template control (NTC) was added to each plate to assess for contamination. MicroAmp Optical Adhesive film (Applied Biosystems, Life Technologies, UK) was used to seal plates. Plates were spun to ensure samples were at the bottom of each well prior to amplification. A StepOnePlus PCR machine (Applied Biosystems, Life Technologies, UK) was used. qPCR runs consisted of an initial 95°C holding stage for 20 seconds, followed by 40 cycles of 95°C (1 second) and 60°C (20 second) and also contained a melt curve step, consisting of 40 cycles of 95°C (15 seconds) and 60°C (1 minute), with a final denaturing step of 95°C (15 seconds).

## Golgi-Cox staining

### Solution make-up

The Golgi-Cox solution was composed of 5% mercuric chloride, 5% potassium dichromate and 5% potassium chromate. 5% mercuric chloride and 5% potassium dichromate were produced by dissolving 10g of mercuric chloride (BDH Chemicals Ltd, Product No. 29163) and 10g of potassium dichromate (BDH Chemicals Ltd, Product No. 0572140), each in 200ml of distilled water. 5% potassium chromate was produced by dissolving 8g of potassium chromate (BDH Chemicals Ltd, Product No. 29598) in 160ml of distilled water. Each dissolving step was performed using a magnetic mixer and heating block. The Golgi-Cox solution was then produced by mixing 200ml of 5% mercuric chloride and 200ml of 5% potassium dichromate and combining with 160ml of 5% potassium chromate mixed with 400ml of distilled water. Once all solutions were mixed, the bottle was covered with aluminium foil and stored in the dark at room temperature for at least 48 hours. A precipitate formed at the bottom of the bottom indicating the correct process had been carried out, which was removed using filter paper before returning the bottle to the previous conditions until usage. All solutions containing mercuric sulfate, potassium chromate and potassium dichromate are toxic, so all actions involving these solutions was carried out with care in a fume cupboard.

Additionally, all waste products were kept in labelled containers under the chemical hood and disposed of via the correct pathways.

The Cryoprotectant solution was formed by dissolving 100g of sucrose and 75ml of glycerol, before making up to 500ml in distilled water.

The 0.1% sodium thiosulphate was formed by dissolving 0.2g of sodium thiosulphate (BDH Chemicals LTD, Product No. 102684G) in 200ml of distilled water.

The 20% ammonia was formed by mixing 20ml ammonia solution (BDH Chemicals LTD, Product No. 10012) with 80ml of distilled water.

### Tissue collection

Following treatment, all rats were anaesthetized by gaseous isoflurane before immediate removal of the head and extraction of the brain. The meninges were carefully removed using tweezers to reduce the likelihood of damaging the brains during the slicing step. To avoid interference with the HPA axis, anesthetizing of the animals was performed in red light, with standard lighting only switched on following removal of the head.

### Golgi-Cox staining procedure

Immediately following removal of the brains, they were placed in falcon tubes containing 15ml of Golgi-Cox solution before storage at room temperature in the dark. The Golgi-Cox solution was then changed once after 24 hours, before a second and third time following week long incubations. The brains were then transferred to falcon tubes containing 20ml of cryoprotectant solution, which was changed daily.

Following at least 3 days in cryoprotectant solution, the brains were sliced into 100μm coronal sections using a vibratome Leica VT1000 S Vibrating blade microtome. Prior to slicing, the speed of the vibrating blade was set as 0.6, the vibration frequency was set as 9 and the vibration amplitude was set as 0.6mm. The brains were stuck to the vibratome base using superglue and carefully oriented perpendicular to the blade as to produce sections matching those shown in The Rat Brain in Stereotaxic Coordinates by Paxinos and Watson. The microtome chamber was then filled with ice-cold PBS and the surrounding water bath filled with water and ice, with the ice being replenished throughout slicing to ensure maintenance of the low temperature. Once cut, floating sections were removed with a brush and transferred into 6 well plates of PBS and stored in the dark at 4 °C.

Once sectioned, the slices were transferred to a large glass container, before being washed for 5 minutes 3 times in distilled water by moving the slices to a fresh container. Next, the slices were transferred to a cell culture dish containing 20% ammonia for 30 minutes at room temperature, before washing twice for 5 minutes as before. The slices were then transferred to a cell culture dish containing 0.1% sodium thiosulfate for 25 minutes in the dark, followed by 2 washes for 5 minutes. Sections were then placed on 1mm thick Superfrost^tm^Plus Adhesion Microscope Slides and semi-dried to help adherence. Care was taken to ensure that the sections did not begin to crack due to more complete drying and the transfer of sections to the slides was done as quickly as possible to allow for even drying across the earlier and later transferred sections.

Once adhered to the slides, dehydration of the sections was carried out by submerging the slides in 50%, 75% and 95% ethanol for 2 minutes each, followed by 100% alcohol for 5 minutes and then xylene for 5 minutes 3 times in 3 different glass containers.

Finally, slides were coverslipped with use of DPX mounting media (CellPath, Reference No. SEA-1300-00A) and stored in the dark at room temperature.

### Histological analysis of Golgi-Cox stained neurons

Brightfield images of stained neurons were captured by a slide scanner (info needed) at both 10x and 40x magnification. Analyzed neurons satisfied the criteria of Layer V pyramidal cells located in the perirhinal cortex that displayed complete staining and were identified using 10x images. 6 neurons were chosen per brain, with random selection occurring in the event that greater than 6 neurons fit the criteria for analysis. Reconstruction of neurons and Sholl analysis was performed using Simple Neurite Tracer, an Fiji software plugin.

Images taken at 40x magnification were used for measurements of spine density, spine length and spine head area, which was done manually using ImageJ measuring tools. Six 20 micron segments were selected per neuron, 3 each for the apical and basal sections of the dendritic tree. Chosen segments were second or greater order dendrites and had no overlap with other dendrites. Apical and basal segments were >50 and >20 microns from the soma, respectively. If more than 3 segments satisfied the criteria for either the apical or basal dendritic of a chosen neuron, 3 segments were chosen at random. Spine density was defined as the number of spines per micron of dendrite length, spine length was the direct distance from the spine head tip to the spine branch point with the shaft and spine head area was the surface area of the head at the tip of each spine. All dendritic analysis was conducted blind to the treatment group of the brains.

### Proteomics

The following protocol was kindly provided by Dr Kate Heesom and the University of Bristol Proteomics facility who carried out the proteomics work presented throughout this work.

### Proteomic analysis

Aliquots of 100µg of eleven samples were digested with trypsin (2.5µg trypsin per 100µg protein; 37°C, overnight), labelled with TMTpro™ 16plex label reagents according to the manufacturer’s protocol (Thermo Fisher Scientific, Loughborough, LE11 5RG, UK) and the labelled samples pooled.

For the Total proteome analysis, an aliquot of 50ug of the pooled sample was desalted using a SepPak cartridge according to the manufacturer’s instructions (Waters, Milford, Massachusetts, USA). Eluate from the SepPak cartridge was evaporated to dryness and resuspended in buffer A (20 mM ammonium hydroxide, pH 10) prior to fractionation by high pH reversed-phase chromatography using an Ultimate 3000 liquid chromatography system (Thermo Fisher Scientific). In brief, the sample was loaded onto an XBridge BEH C18 Column (130Å, 3.5 µm, 2.1 mm X 150 mm, Waters, UK) in buffer A and peptides eluted with an increasing gradient of buffer B (20 mM Ammonium Hydroxide in acetonitrile, pH 10) from 0-95% over 60 minutes. The resulting fractions were evaporated to dryness and resuspended in 1% formic acid prior to analysis by nano-LC MSMS using an Orbitrap Fusion Lumos mass spectrometer (Thermo Scientific).

High pH RP fractions (Total proteome analysis) or the phospho-enriched fractions (Phospho-proteome analysis) were further fractionated using an Ultimate 3000 nano-LC system in line with an Orbitrap Fusion Lumos mass spectrometer (Thermo Scientific). In brief, peptides in 1% (vol/vol) formic acid were injected onto an Acclaim PepMap C18 nano-trap column (Thermo Scientific). After washing with 0.5% (vol/vol) acetonitrile 0.1% (vol/vol) formic acid peptides were resolved on a 250 mm × 75 μm Acclaim PepMap C18 reverse phase analytical column (Thermo Scientific) over a 150 min organic gradient, using 7 gradient segments (1-6% solvent B over 1min., 6-15% B over 58min., 15-32%B over 58min., 32-40%B over 5min., 40-90%B over 1min., held at 90%B for 6min and then reduced to 1%B over 1min.) with a flow rate of 300 nl min−1. Solvent A was 0.1% formic acid and Solvent B was aqueous 80% acetonitrile in 0.1% formic acid. Peptides were ionized by nano-electrospray ionization at 2.0kV using a stainless-steel emitter with an internal diameter of 30 μm (Thermo Scientific) and a capillary temperature of 300°C.

All spectra were acquired using an Orbitrap Fusion Lumos mass spectrometer controlled by Xcalibur 3.0 software (Thermo Scientific) and operated in data-dependent acquisition mode using an SPS-MS3 workflow. FTMS1 spectra were collected at a resolution of 120 000, with an automatic gain control (AGC) target of 200 000 and a max injection time of 50ms. Precursors were filtered with an intensity threshold of 5000, according to charge state (to include charge states 2-7) and with monoisotopic peak determination set to Peptide. Previously interrogated precursors were excluded using a dynamic window (60s +/-10ppm). The MS2 precursors were isolated with a quadrupole isolation window of 0.7m/z. ITMS2 spectra were collected with an AGC target of 10 000, max injection time of 70ms and CID collision energy of 35%.

For FTMS3 analysis, the Orbitrap was operated at 50 000 resolution with an AGC target of 50 000 and a max injection time of 105ms. Precursors were fragmented by high energy collision dissociation (HCD) at a normalised collision energy of 60% to ensure maximal TMT reporter ion yield. Synchronous Precursor Selection (SPS) was enabled to include up to 10 MS2 fragment ions in the FTMS3 scan.

For the Phospho proteome analysis, the remainder of the TMT-labelled pooled sample was also desalted using a SepPak cartridge (Waters, Milford, Massachusetts, USA).

Eluate from the SepPak cartridge was evaporated to dryness and subjected to TiO2-based phosphopeptide enrichment according to the manufacturer’s instructions (Pierce). The flow-through and washes from the TiO2-based enrichment were then subjected to FeNTA-based phosphopeptide enrichment according to the manufacturer’s instructions (Pierce). The phospho-enriched samples were again evaporated to dryness and then resuspended in 1% formic acid prior to analysis by nano-LC MSMS using an Orbitrap Fusion Lumos mass spectrometer (Thermo Scientific).

### Nano-LC Mass Spectrometry

High pH RP fractions (Total proteome analysis) or the phospho-enriched fractions (Phospho-proteome analysis) were further fractionated using an Ultimate 3000 nano-LC system in line with an Orbitrap Fusion Lumos mass spectrometer (Thermo Scientific). In brief, peptides in 1% (vol/vol) formic acid were injected onto an Acclaim PepMap C18 nano-trap column (Thermo Scientific). After washing with 0.5% (vol/vol) acetonitrile 0.1% (vol/vol) formic acid peptides were resolved on a 250 mm × 75 μm Acclaim PepMap C18 reverse phase analytical column (Thermo Scientific) over a 150 min organic gradient, using 7 gradient segments (1-6% solvent B over 1min., 6-15% B over 58min., 15-32%B over 58min., 32-40%B over 5min., 40-90%B over 1min., held at 90%B for 6min and then reduced to 1%B over 1min.) with a flow rate of 300 nl min−1. Solvent A was 0.1% formic acid and Solvent B was aqueous 80% acetonitrile in 0.1% formic acid. Peptides were ionized by nano-electrospray ionization at 2.0kV using a stainless-steel emitter with an internal diameter of 30 μm (Thermo Scientific) and a capillary temperature of 300°C.

All spectra were acquired using an Orbitrap Fusion Lumos mass spectrometer controlled by Xcalibur 3.0 software (Thermo Scientific) and operated in data-dependent acquisition mode using an SPS-MS3 workflow. FTMS1 spectra were collected at a resolution of 120 000, with an automatic gain control (AGC) target of 200 000 and a max injection time of 50ms. Precursors were filtered with an intensity threshold of 5000, according to charge state (to include charge states 2-7) and with monoisotopic peak determination set to Peptide. Previously interrogated precursors were excluded using a dynamic window (60s +/-10ppm). The MS2 precursors were isolated with a quadrupole isolation window of 0.7m/z. ITMS2 spectra were collected with an AGC target of 10 000, max injection time of 70ms and CID collision energy of 35%.

For FTMS3 analysis, the Orbitrap was operated at 50 000 resolution with an AGC target of 50 000 and a max injection time of 105ms. Precursors were fragmented by high energy collision dissociation (HCD) at a normalised collision energy of 60% to ensure maximal TMT reporter ion yield. Synchronous Precursor Selection (SPS) was enabled to include up to 10 MS2 fragment ions in the FTMS3 scan.

## Data Analysis

The raw data files were processed and quantified using Proteome Discoverer software v2.4 (Thermo Scientific) and searched against the UniProt Rattus Norvegicus canonical sequences database (downloaded February 2023: 47939 entries) using the SEQUEST HT algorithm. Peptide precursor mass tolerance was set at 10ppm, and MS/MS tolerance was set at 0.6Da. Search criteria included oxidation of methionine (+15.995Da), acetylation of the protein N-terminus (+42.011Da) and methionine loss plus acetylation of the protein N-terminus (-89.03Da) as variable modifications and carbamidomethylation of cysteine (+57.0214) and the addition of the TMTpro™ mass tag (+304.2071) to peptide N-termini and lysine as fixed modifications. For the Phospho-proteome analysis, phosphorylation of serine, threonine and tyrosine (+79.966) was also included as a variable modification. Searches were performed with full tryptic digestion and a maximum of 2 missed cleavages were allowed. The reverse database search option was enabled and all data was filtered to satisfy false discovery rate (FDR) of 5%.

## Pathway analysis

The pathway analysis included in this work was carried out by Dr Phil Lewis. Raw data composing of the master protein accessions, phospho-sites, and associated p-values, FDR-adjusted p-values and Log2 fold-changes were uploaded to QIAGEN Ingenuity Pathway Analysis (IPA) software. Both total protein and phosphorylation analyses were performed with a p-value cut-off of p<0.05 to allow canonical pathway analysis to identify the pathways from the QIAGEN Ingenuity Pathway Analysis library of canonical pathways that saw significant dysregulation in the inputted raw dataset.

## Brain slice preparation

Adult male rats (230-300 g) were anaesthetized with isoflurane by inhalation and killed by decapitation. The brain was then swiftly removed and placed in to ice cold modified sucrose artificial cerebrospinal fluid (aCSF) (containing (in mM): 52.5 NaCl, 2.5 KCl, 25 NaHCO_3_, 1.25 NaH_2_PO_4_, 5 MgCl_2_, 25 D-Glucose, 100 Sucrose, 2 CaCl_2_, 0.1 Kynurenic acid) that had been saturated with 95% O_2_ and 5% CO_2_. The brain was removed from solution and quickly blocked on filter paper on a glass petri dish filled with ice. The brain was hemisected along the sagittal midline and a 45° cut was made from the caudal end to the dorsal edge of each hemisphere. The halves were then glued (cyanoacrylate adhesive) by this cut side to a metal vibratome stage and supported with a piece of 2% w/v agar.

The brain was immediately submerged in ice cold carbogenated modified sucrose aCSF within the slicing chamber and 300 or 400 μm parasagittal slices cut using a vibratome (Leica VT1000S). Slices were transferred to a petri dish containing room temperature (RT) aCSF (containing (in mM): 124 NaCl, 3 KCl, 26 NaHCO_3_, 1.4 NaH_2_PO_4_, 1 MgSO4, 10 D-Glucose, 2 CaCl_2_), wherein the perirhinal slice was isolated and excess tissue discarded. Perirhinal slices were then transferred to the holding chamber filled with fresh aCSF and incubated at 32 °C for 30 minutes. The slices were then held for > 30 minutes at RT before electrophysiological recording.

## Field Electrophysiological recordings

Slices were transferred to a submersion style recording chamber maintained between 28-30 °C and constantly perfused with aCSF saturated with carbogen at a rate of ∼2 ml/min. Electrical signals were recorded using glass capillaries pulled using a microelectrode puller (Sutter Instruments) at a resistance of 2-6 MΩ, filled with aCSF. Signals were amplified using an AxoClamp 2B amplifier (Axon Instruments) and 50/60 Hz noise eliminated by a Hum Bug (Quest Scientific), data was acquired at 10 KHz and analysed using WinLTP software[77]. Field excitatory post-synaptic potentials (fEPSPs) were evoked at 0.033 Hertz (Hz) by placing bipolar stimulating electrodes on either side of the recording electrode which was positioned directly beneath the rhinal sulcus at the juxtaposition of layer I and layers II/III. One stimulating electrode was placed superficial and one deep to ensure independent responses. A stable baseline recording of 30 minutes was acquired before experimental manipulations were applied.

Compounds were applied by addition to the perfusing aCSF.

## Whole cell patch clamp recordings

Slices were transferred into a submersion chamber as described above and constantly perfused with aCSF. Microelectrodes (3-6 MΩ) were filled with intracellular solution (in mM: 120 CsMeSO3, 5 CsCl, 2.8 NaCl, 3 MgCl2, 20 HEPES, 5 EGTA, 0.5 CaCl2, 2 Adenosine 5′-triphosphate magnesium salt, 0.3 Guanosine 5′-triphosphate sodium salt hydrate 5 QX-314) with pH adjusted to 7.25 using CsOH. Osmolarity of 295 mOsmoles/kg was achieved and solution passed through a 0.2 μm filter before patching. A bipolar stimulating electrode was placed to elicit action potentials at 0.07 Hz in fibres of layers II/III and whole cell recordings were taken from pyramidal neurons. Series resistance was monitored throughout the course of the experiment and experiments with changes of >25% were discarded.

## Novel Object Recognition (NOR) memory task

For all memory testing, different rats were used at all time points and treatment. Rats were transferred to a sound-attenuating behavior room in low light (40 to 50 lx on arena floor) at ZT11. Rats were left to habituate to the room for at least an hour, prior to starting behavioral experiments. Rats were handled for 1 week in the behavior room prior to experimental start, followed by habituation to the arena without stimuli for 10 min daily for 5 d. Training and testing occurred in an open-top arena (50 × 90 × 100 cm) made of wood. The walls inside the arena were black and floor covered in sawdust. An overhead camera recorded behavior for analysis. Exploration was scored when the rat head orientated toward the object and came within 1 cm of the object. The objects were constructed from Duplo blocks, which were too heavy for the animals to displace.

During training, rats were exposed to two identical objects and allowed to explore for 4 min. These objects were placed in the far side of the arena, 10 cm away from the walls, to allow full access around the objects. During the retention test (1 h for short-term memory, 6 h for intermediate memory, or 24 h for long-term memory), rats were allowed to explore for 3 min. Prior to testing, one of the objects was replaced with a novel object discernible in colour and shape from the previous object. The position of the object was counterbalanced between rats. Total exploration time was recorded, and preference for the novel item was expressed as a discrimination index.

## References

1. Herman, J.P. and W.E. Cullinan, Neurocircuitry of stress: central control of the hypothalamo– pituitary–adrenocortical axis. Trends in neurosciences, 1997. 20(2): p. 78–84.

2. Kalsbeek, A., et al., Circadian rhythms in the hypothalamo–pituitary–adrenal (HPA) axis. Molecular and cellular endocrinology, 2012. 349(1): p. 20–29.

3. Exton, J., Regulation of gluconeogenesis by glucocorticoids. Monographs on endocrinology, 1979. 12: p. 535–546.

4. Schleimer, R., An overview of glucocorticoid anti-inflammatory actions. European journal of clinical pharmacology, 1993. 45: p. S3–S7.

5. Cain, D.W. and J.A. Cidlowski, Immune regulation by glucocorticoids. Nature Reviews Immunology, 2017. 17(4): p. 233–247.

6. Pariante, C.M. and S.L. Lightman, The HPA axis in major depression: classical theories and new developments. Trends in neurosciences, 2008. 31(9): p. 464–468.

7. Tsui, A., et al., Longitudinal associations between diurnal cortisol variation and later-life cognitive impairment. Neurology, 2020. 94(2): p. e133–e141.

8. Amasi-Hartoonian, N., et al., Cause or consequence? Understanding the role of cortisol in the increased inflammation observed in depression. Current opinion in endocrine and metabolic research, 2022. 24: p. 100356.

9. Birnie, M.T., et al., Circadian regulation of hippocampal function is disrupted with corticosteroid treatment. Proceedings of the National Academy of Sciences, 2023. 120(15): p. e2211996120.

10. Liston, C., et al., Circadian glucocorticoid oscillations promote learning-dependent synapse formation and maintenance. Nature neuroscience, 2013. 16(6): p. 698.

11. Karst, H., et al., Metaplasticity of amygdalar responses to the stress hormone corticosterone. Proceedings of the National Academy of Sciences, 2010. 107(32): p. 14449–14454.

12. den Boon, F.S., et al., Circadian and ultradian variations in corticosterone level influence functioning of the male mouse basolateral amygdala. Endocrinology, 2019. 160(4): p. 791–802.

13. Sarabdjitsingh, R.A., et al., Ultradian corticosterone pulses balance glutamatergic transmission and synaptic plasticity. Proceedings of the National Academy of Sciences, 2014. 111(39): p. 14265–14270.

14. Sarabdjitsingh, R.A., et al., Hippocampal fast glutamatergic transmission is transiently regulated by corticosterone pulsatility. PloS one, 2016. 11(1).

15. Buckley, M.J. and D. Gaffan, Perirhinal cortical contributions to object perception. Trends in cognitive sciences, 2006. 10(3): p. 100–107.

16. Brown, M.W. and J.P. Aggleton, Recognition memory: what are the roles of the perirhinal cortex and hippocampus? Nature Reviews Neuroscience, 2001. 2(1): p. 51–61.

17. Suzuki, W.L. and D.G. Amaral, Perirhinal and parahippocampal cortices of the macaque monkey: cortical afferents. Journal of comparative neurology, 1994. 350(4): p. 497–533.

18. De Curtis, M. and D. Paré, The rhinal cortices: a wall of inhibition between the neocortex and the hippocampus. Progress in neurobiology, 2004. 74(2): p. 101–110.

19. Koganezawa, N., et al., Significance of the deep layers of entorhinal cortex for transfer of both perirhinal and amygdala inputs to the hippocampus. Neuroscience research, 2008. 61(2): p. 172–181.

20. Burwell, R.D., M.P. Witter, and D.G. Amaral, Perirhinal and postrhinal cortices of the rat: a review of the neuroanatomical literature and comparison with findings from the monkey brain. Hippocampus, 1995. 5(5): p. 390–408.

21. Squire, L.R., C.E. Stark, and R.E. Clark, The medial temporal lobe. Annu. Rev. Neurosci., 2004. 27: p. 279–306.

22. Aggleton, J.P., R.J. Kyd, and D.K. Bilkey, When is the perirhinal cortex necessary for the performance of spatial memory tasks? Neuroscience & Biobehavioral Reviews, 2004. 28(6): p. 611–624.

23. Zhu, X., et al., Effects of the novelty or familiarity of visual stimuli on the expression of the immediate early gene c-fos in rat brain. Neuroscience, 1995. 69(3): p. 821–829.

24. Lightman, S.L. and B.L. Conway-Campbell, The crucial role of pulsatile activity of the HPA axis for continuous dynamic equilibration. Nature Reviews Neuroscience, 2010. 11(10): p. 710.

25. Stavreva, D.A., et al., Ultradian hormone stimulation induces glucocorticoid receptor-mediated pulses of gene transcription. Nature cell biology, 2009. 11(9): p. 1093–1102.

26. Kalafatakis, K., et al., Ultradian rhythmicity of plasma cortisol is necessary for normal emotional and cognitive responses in man. Proceedings of the National Academy of Sciences, 2018. 115(17): p. E4091–E4100.

27. Christoffel, D.J., et al., IκB kinase regulates social defeat stress-induced synaptic and behavioral plasticity. Journal of Neuroscience, 2011. 31(1): p. 314–321.

28. Aleisa, A.M., et al., Nicotine blocks stress-induced impairment of spatial memory and long-term potentiation of the hippocampal CA1 region. The The International Journal of Neuropsychopharmacology, 2006. 9(4): p. 417–426.

29. Rich, M.T., et al., Phosphoproteomic analysis reveals a novel mechanism of CaMKIIα regulation inversely induced by cocaine memory extinction versus reconsolidation. Journal of Neuroscience, 2016. 36(29): p. 7613–7627.

30. Wang, P., et al., PTENα modulates CaMKII signaling and controls contextual fear memory and spatial learning. Cell reports, 2017. 19(12): p. 2627–2641.

31. Klann, E. and T.E. Dever, Biochemical mechanisms for translational regulation in synaptic plasticity. Nature Reviews Neuroscience, 2004. 5(12): p. 931–942.

32. Zhou, S. and Y. Yu, Synaptic EI balance underlies efficient neural coding. Frontiers in neuroscience, 2018. 12: p. 46.

33. Shew, W.L., et al., Information capacity and transmission are maximized in balanced cortical networks with neuronal avalanches. Journal of neuroscience, 2011. 31(1): p. 55–63.

34. Sohal, V.S. and J.L. Rubenstein, Excitation-inhibition balance as a framework for investigating mechanisms in neuropsychiatric disorders. Molecular psychiatry, 2019. 24(9): p. 1248–1257.

35. Radley, J.J., et al., Repeated stress alters dendritic spine morphology in the rat medial prefrontal cortex. J Comp Neurol, 2008. 507(1): p. 1141–50.

36. Radley, J.J., et al., Repeated stress induces dendritic spine loss in the rat medial prefrontal cortex. Cereb Cortex, 2006. 16(3): p. 313–20.

37. Duprat, F., et al., GluR2 protein-protein interactions and the regulation of AMPA receptors during synaptic plasticity. Philosophical Transactions of the Royal Society of London Series B-Biological Sciences, 2003. 358(1432): p. 715–720.

38. Franchini, L., et al., Synaptic GluN2A-Containing NMDA Receptors: From Physiology to Pathological Synaptic Plasticity. International Journal of Molecular Sciences, 2020. 21(4).

39. Ashby, M.C., et al., Removal of AMPA receptors (AMPARs) from synapses is preceded by transient endocytosis of extrasynaptic AMPARs. Journal of Neuroscience, 2004. 24(22): p. 5172–5176.

40. Lee, S., et al., SHP2 regulates GluA2 tyrosine phosphorylation required for AMPA receptor endocytosis and mGluR-LTD. Proceedings of the National Academy of Sciences of the United States of America, 2024. 121(18).

41. McCormack, S.G., R.L. Stornetta, and J.J. Zhu, Synaptic AMPA receptor exchange maintains bidirectional plasticity. Neuron, 2006. 50(1): p. 75–88.

42. Ma, H., et al., Amygdala-hippocampal innervation modulates stress-induced depressive-like behaviors through AMPA receptors. Proceedings of the National Academy of Sciences of the United States of America, 2021. 118(6).

43. Griffiths, S., et al., Expression of long-term depression underlies visual recognition memory. Neuron, 2008. 58(2): p. 186–194.

44. Massey, P.V., et al., Differential roles of NR2A and NR2B-containing NMDA receptors in cortical long-term potentiation and long-term depression. Journal of Neuroscience, 2004. 24(36): p. 7821–7828.

45. Flynn, B.P., B.L. Conway-Campbell, and S.L. Lightman, The emerging importance of ultradian glucocorticoid rhythms within metabolic pathology. Ann Endocrinol (Paris), 2018. 79(3): p. 112–114.

46. Claydon, M.D. and B.L. Conway–Campbell, The glucocorticoid-mediated genomic stress response. Current Opinion in Endocrine and Metabolic Research, 2022. 25: p. 100363.

47. Cho, K., et al., A new form of long-term depression in the perirhinal cortex. Nature neuroscience, 2000. 3(2): p. 150–156.

48. Ziakopoulos, Z., et al., Input-and layer-dependent synaptic plasticity in the rat perirhinal cortex in vitro. Neuroscience, 1999. 92(2): p. 459–472.

49. Bliss, T.V. and G.L. Collingridge, A synaptic model of memory: long-term potentiation in the hippocampus. Nature, 1993. 361(6407): p. 31–39.

50. Bogacz, R. and M.W. Brown, An anti-Hebbian model of familiarity discrimination in the perirhinal cortex. Neurocomputing, 2003. 52: p. 1–6.

51. Devasani, K. and Y. Yao, Expression and functions of adenylyl cyclases in the CNS. Fluids and Barriers of the CNS, 2022. 19(1): p. 23.

52. Villacres, E.C., et al., Type I adenylyl cyclase mutant mice have impaired mossy fiber long-term potentiation. Journal of Neuroscience, 1998. 18(9): p. 3186–3194.

53. Zhang, M. and H. Wang, Ca2+-stimulated ADCY1 and ADCY8 regulate distinct aspects of synaptic and cognitive flexibility. Frontiers in Cellular Neuroscience, 2023. 17: p. 1215255.

54. Wang, H., et al., Type 8 adenylyl cyclase is targeted to excitatory synapses and required for mossy fiber long-term potentiation. Journal of Neuroscience, 2003. 23(30): p. 9710–9718.

55. Gilabert-Juan, J., et al., Reduced interneuronal dendritic arborization in CA1 but not in CA3 region of mice subjected to chronic mild stress. Brain Behav, 2017. 7(2): p. e00534.

56. Vyas, A., et al., Chronic stress induces contrasting patterns of dendritic remodeling in hippocampal and amygdaloid neurons. J Neurosci, 2002. 22(15): p. 6810–8.

57. Lakshminarasimhan, H. and S. Chattarji, Stress Leads to Contrasting Effects on the Levels of Brain Derived Neurotrophic Factor in the Hippocampus and Amygdala. Plos One, 2012. 7(1).

58. Herms, J. and M.M. Dorostkar, Dendritic Spine Pathology in Neurodegenerative Diseases. Annu Rev Pathol, 2016. 11: p. 221–50.

59. Cerqueira, J.J., et al., Specific configuration of dendritic degeneration in pyramidal neurons of the medial prefrontal cortex induced by differing corticosteroid regimens. Cereb Cortex, 2007. 17(9): p. 1998–2006.

60. Silva-Gomez, A.B., et al., Dexamethasone induces different morphological changes in the dorsal and ventral hippocampus of rats. J Chem Neuroanat, 2013. 47: p. 71–8.

61. Ge, M.J., et al., Chronic restraint stress induces depression-like behaviors and alterations in the afferent projections of medial prefrontal cortex from multiple brain regions in mice. Brain Research Bulletin, 2024. 213.

62. Kasai, H., et al., Structure-stability-function relationships of dendritic spines. Trends Neurosci, 2003. 26(7): p. 360–8.

63. Bronk, P., et al., Differential effects of SNAP-25 deletion on Ca2+-dependent and Ca2+-independent neurotransmission. Journal of neurophysiology, 2007. 98(2): p. 794–806.

64. Washbourne, P., et al., Genetic ablation of the t-SNARE SNAP-25 distinguishes mechanisms of neuroexocytosis. Nature neuroscience, 2002. 5(1): p. 19–26.

65. Tafoya, L.C., et al., Expression and function of SNAP-25 as a universal SNARE component in GABAergic neurons. Journal of Neuroscience, 2006. 26(30): p. 7826–7838.

66. Warburton, E.C., et al., Cholinergic neurotransmission is essential for perirhinal cortical plasticity and recognition memory. Neuron, 2003. 38(6): p. 987–996.

67. Barker, G.R.I., et al., A temporally distinct role for group I and group II metabotropic glutamate receptors in object recognition memory. Learning & Memory, 2006. 13(2): p. 178–186.

68. Xiang, J.-Z. and M. Brown, Differential neuronal encoding of novelty, familiarity and recency in regions of the anterior temporal lobe. Neuropharmacology, 1998. 37(4-5): p. 657–676.

69. Barker, G.R., et al., The different effects on recognition memory of perirhinal kainate and NMDA glutamate receptor antagonism: implications for underlying plasticity mechanisms. Journal of Neuroscience, 2006. 26(13): p. 3561–3566.

70. Winters, B.D. and T.J. Bussey, Glutamate receptors in perirhinal cortex mediate encoding, retrieval, and consolidation of object recognition memory. Journal of Neuroscience, 2005. 25(17): p. 4243–4251.

71. Torrisi, S.A., et al., Acute stress alters recognition memory and AMPA/NMDA receptor subunits in a sex-dependent manner. Neurobiol Stress, 2023. 25: p. 100545.

72. Luine, V., et al., Sex differences in chronic stress effects on cognition in rodents. Pharmacol Biochem Behav, 2017. 152: p. 13–19.

73. Vierk, R., et al., Aromatase inhibition abolishes LTP generation in female but not in male mice. Journal of Neuroscience, 2012. 32(24): p. 8116–8126.

74. Wang, W., et al., Memory-related synaptic plasticity is sexually dimorphic in rodent hippocampus. Journal of Neuroscience, 2018. 38(37): p. 7935–7951.

75. Keenan, P., et al., The effect on memory of chronic prednisone treatment in patients with systemic disease. Neurology, 1996. 47(6): p. 1396–1402.

76. Cuestas Torres, D.M. and F.P. Cardenas, Synaptic plasticity in Alzheimer’s disease and healthy aging. Reviews in the Neurosciences, 2020. 31(3): p. 245–268.

77. Anderson, W.W. and G.L. Collingridge, Capabilities of the WinLTP data acquisition program extending beyond basic LTP experimental functions. Journal of neuroscience methods, 2007. 162(1-2): p. 346–356.

