## Supplementary material for "Perirhinal cortex structure and function is dysregulated by corticosteroid treatment"

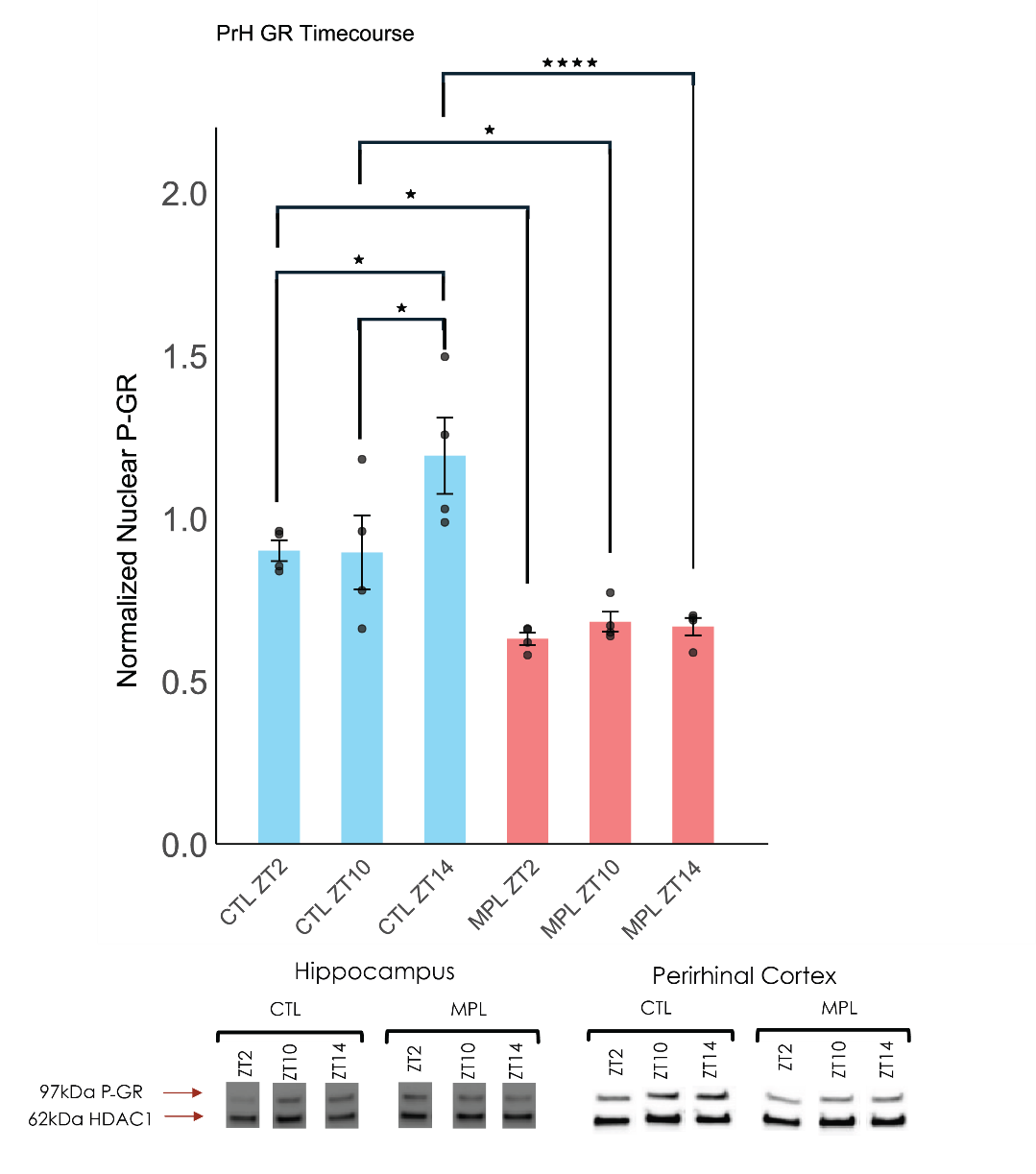

Figure S1. Nuclear GR is suppressed by MPL treatment throughout the circadian cycle in the perirhinal cortex

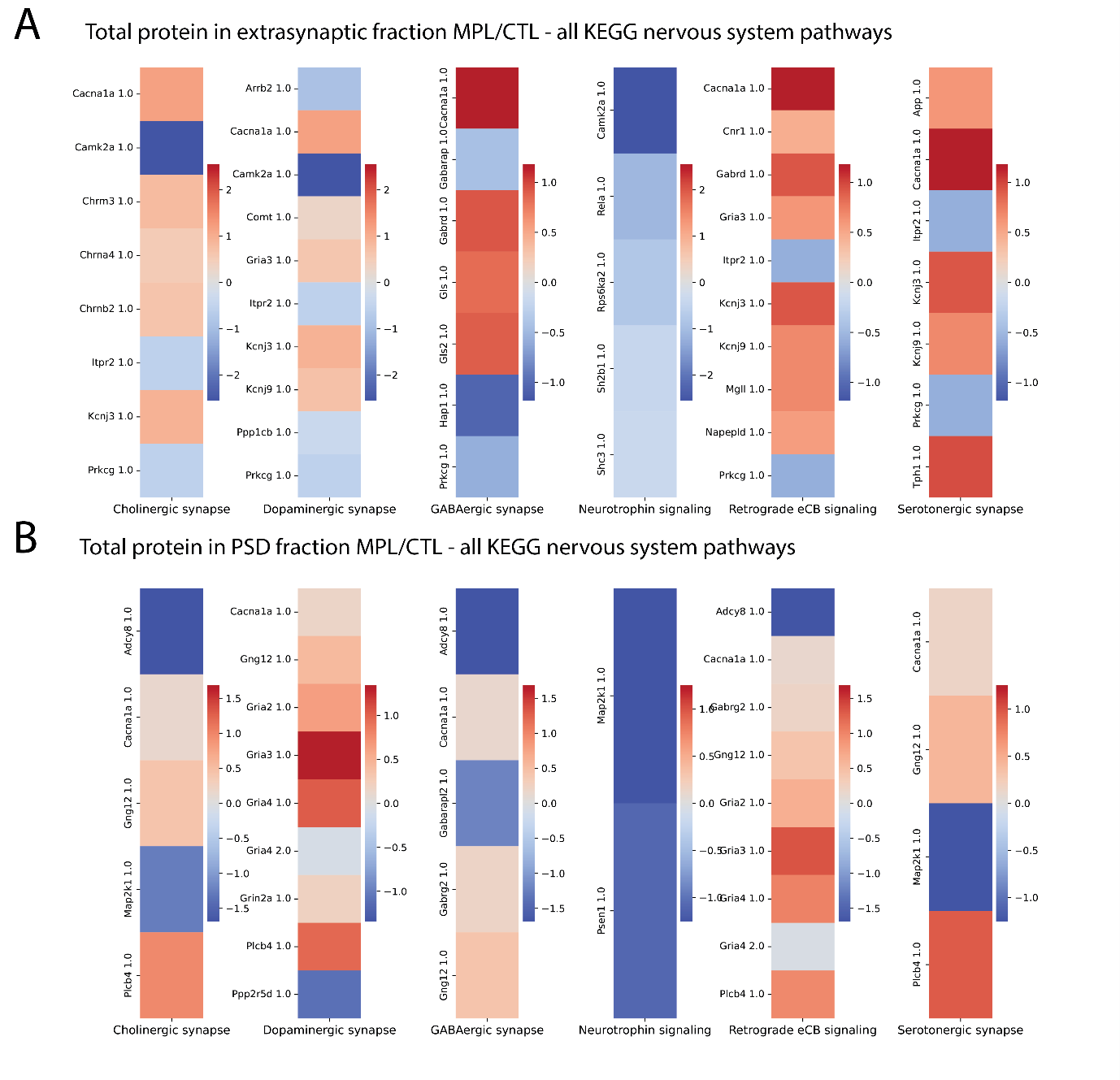

Figure S2. Quantification of fold change of proteins expressed associated with nervous system pathways according to the Kyoto Encyclopaedia of Genes and Genomes (KEGG).

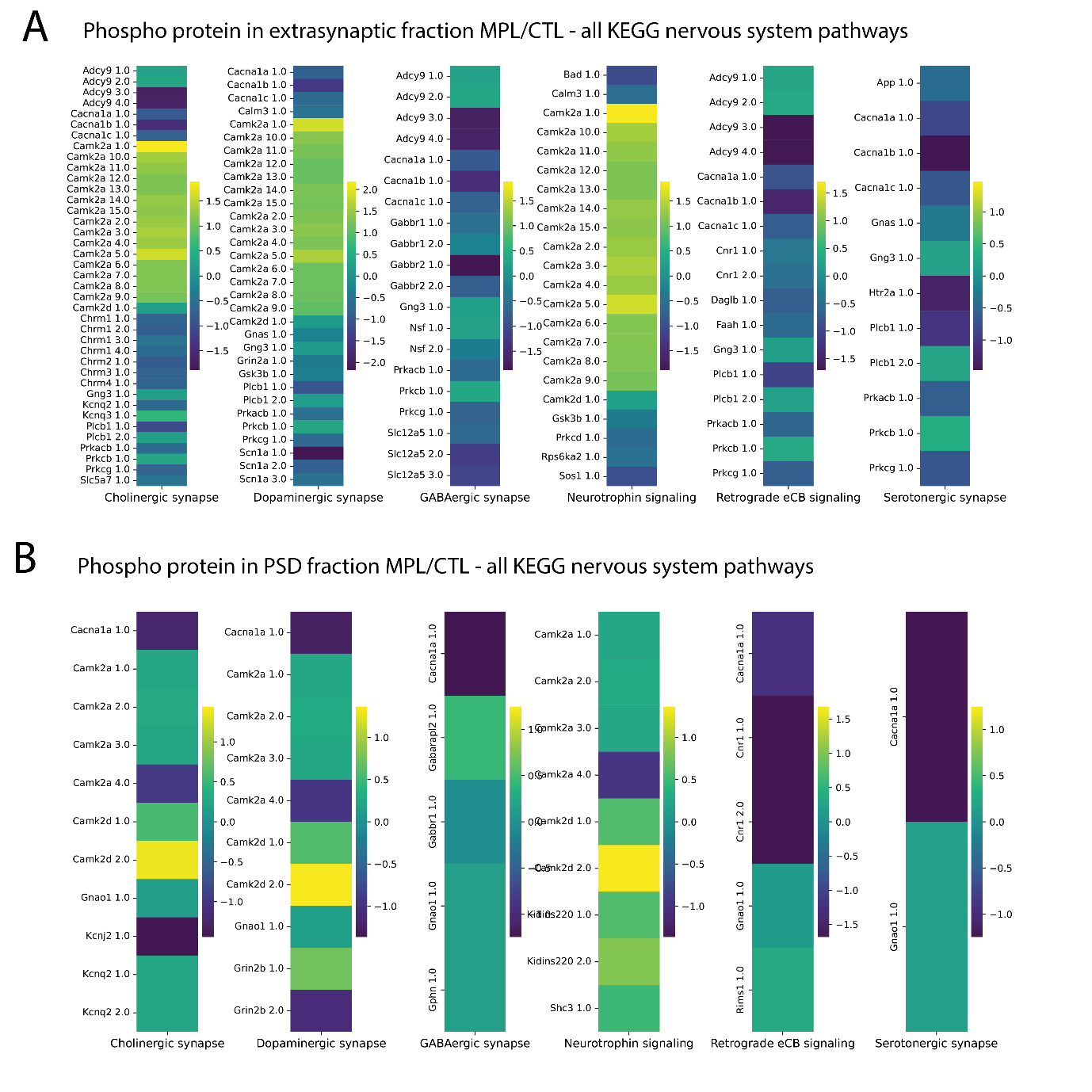

Figure S3. Quantification of changes to phospho-protein associated with other KEGG nervous system pathways.

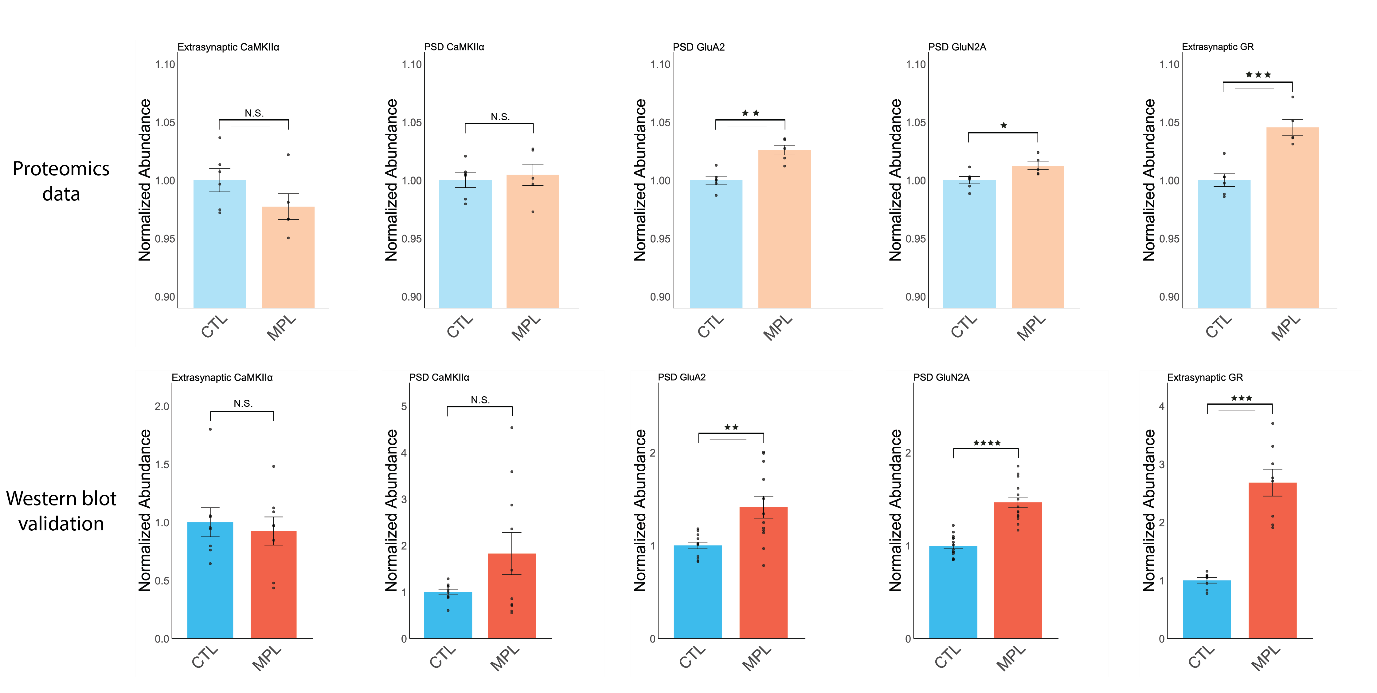

Figure S4. Western blot validation of key protein changes identified in the proteomics dataset

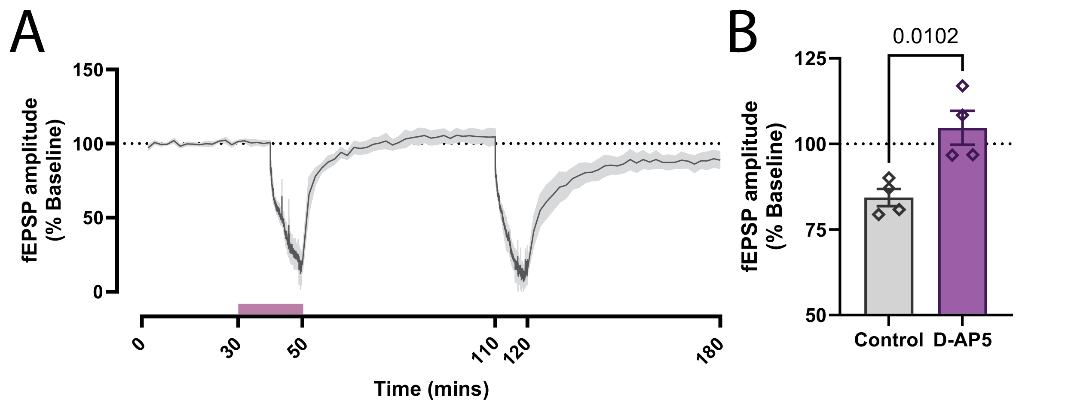

Figure S5. Perirhinal cortex LTD is dependent on NMDA receptor activation

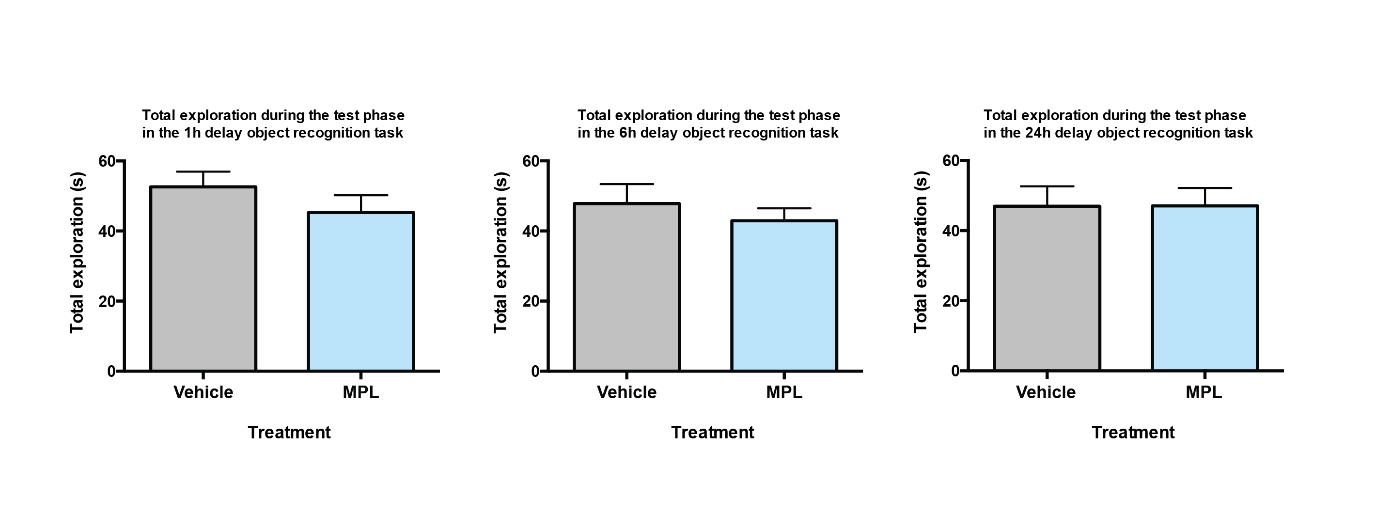

Figure S6. Total exploration in the NOR task was unchanged at all sample-test intervals

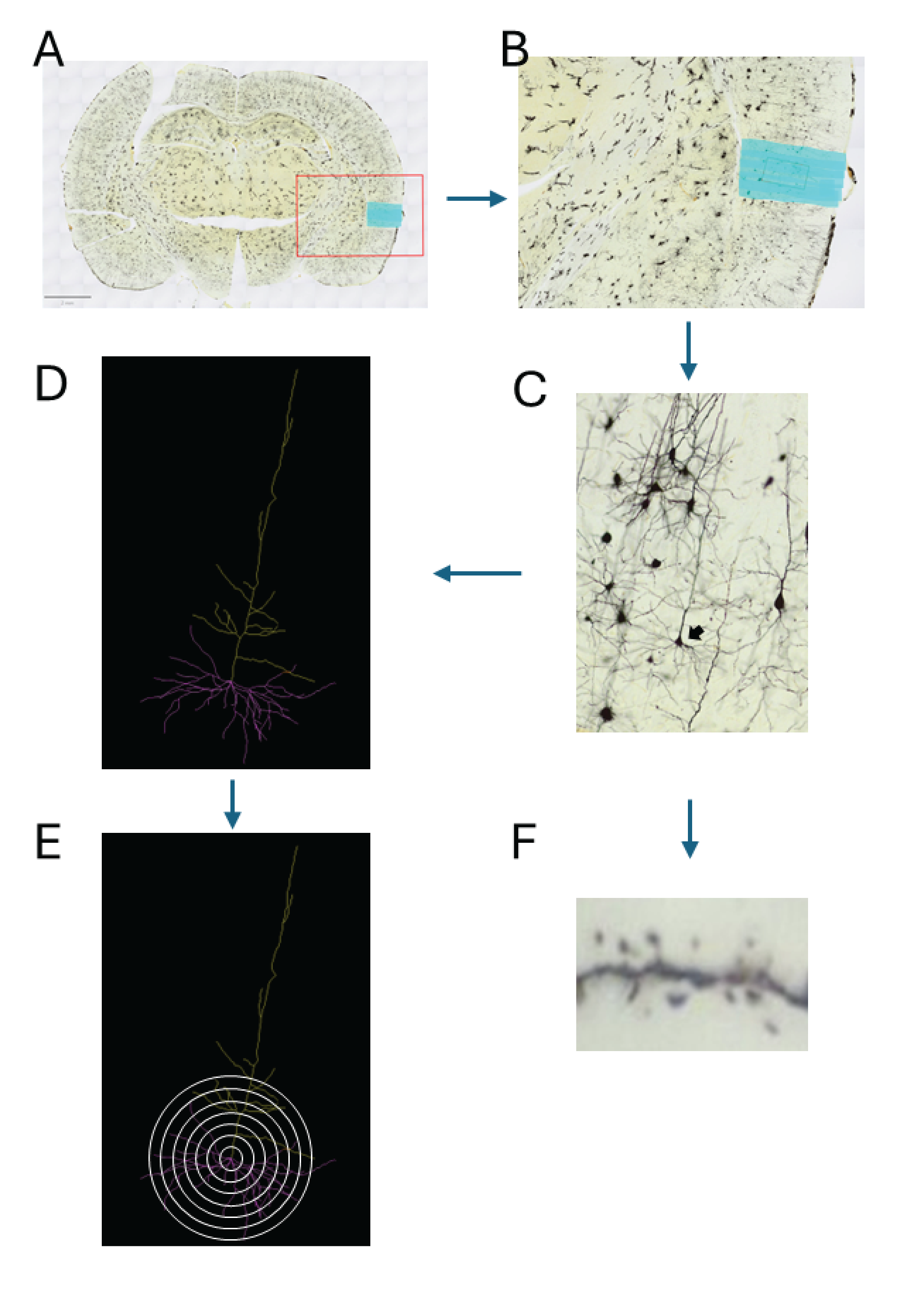

Figure S7. Golgi cox staining followed by Scholl and spine analyses

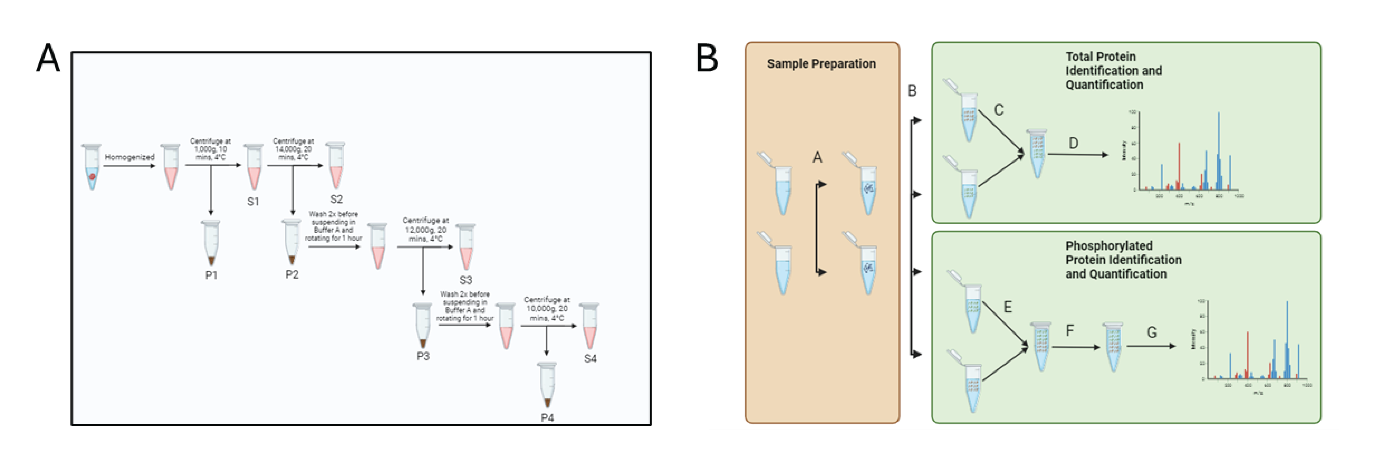

Figure S8. Synaptic fractionation protocol for the preparation of extra-synaptic and post-synaptic density fractions for proteomics.
